# Lactobacilli and other gastrointestinal microbiota of *Peromyscus leucopus*, reservoir host for agents of Lyme disease and other zoonoses in North America

**DOI:** 10.1101/2020.04.02.021659

**Authors:** Ana Milovic, Khalil Bassam, Hanjuan Shao, Ioulia Chatzistamou, Danielle M. Tufts, Maria Diuk-Wasser, Alan G. Barbour

## Abstract

The cricetine rodent *Peromyscus leucopus* is an important reservoir for several human zoonoses, including Lyme disease, in North America. Akin to hamsters, the white-footed deermouse has been unevenly characterized in comparison to the murid *Mus musculus*. To further understanding of *P. leucopus*’ total genomic content, we investigated gut microbiomes of an outbred colony of *P. leucopus*, inbred *M. musculus*, and a natural population of *P. leucopus*. Metagenome and whole genome sequencing were combined with microbiology and microscopy approaches. A focus was the genus *Lactobacillus*, four diverse species of which were isolated from forestomach and feces of colony *P. leucopus*. Three of the species--*L. animalis*, *L. reuteri*, and provisionally-named species “L. peromysci”--were identified in fecal metagenomes of wild *P. leucopus* but not discernibly in samples from *M. musculus*. *L. johnsonii*, the fourth species, was common in *M. musculus* but absent or sparse in wild *P. leucopus*. Also identified in both colony and natural populations were a *Helicobacter* sp. in feces but not stomach, and a *Tritrichomonas* sp. protozoan in cecum or feces. The gut metagenomes of colony *P. leucopus* were similar to those of colony *M. musculus* at the family or higher level and for major subsystems. But there were multiple differences between species and sexes within each species in their gut metagenomes at orthologous gene level. These findings provide a foundation for hypothesis-testing of functions of individual microbial species and for interventions, such as bait vaccines based on an autochthonous bacterium and targeting *P. leucopus* for transmission-blocking.

## Introduction

Epigraph: “I have always looked at problems from an ecological point of view, by placing most emphasis not on the living things themselves, but rather on their inter-relationships and on their interplay with surroundings and events.” René Dubos, 1981 (1, 2)

*Peromyscus leucopus*, the white-footed deermouse, is one of the most abundant wild mammals in central and eastern United States and adjacent regions of Canada and Mexico (3, 4). The rodent is an omnivore, consuming a variety of seeds, such as oak acorns, as well as insects and other invertebrates. Its wide geographic range extends from rural areas to suburbs and even cities, and it is especially common in areas where humans and wildland areas interface (5). Conditions permitting, *P. leucopus* is procreatively proliferant, with 20 or more litters during a female’s period of fecundity (6).

Although commonly called a “mouse”, this species and other members of the genus *Peromyscus* belong to the family Cricetidae, which includes hamsters and voles, and not the family Muridae, which includes *Mus* and *Rattus*. The pairwise divergence time for the genera *Peromyscus* and *Mus* is estimated to be ∼27 million years ago (7, 8), approximately the time since divergence of the family Hominidae, the great apes and hominids, from Cercopithecidae, the Old World monkeys (7, 9). While only a minority of a birth cohort of *P. leucopus* typically survive the predation and winter conditions of their first year in nature (10, 11), in captivity *Peromyscus* species can live twice as long as the laboratory mouse or rat (12). *P. leucopus* differs in its social behavior and reproductive physiology from rodents that are traditional experimental models (13, 14).

*P. leucopus* also merits special attention as a natural host and keystone reservoir for several tickborne zoonoses of humans (reviewed in (15). These include Lyme disease, babesiosis, anaplasmosis, a form of relapsing ever, an ehrlichiosis, and a viral encephalitis. For humans these infections are commonly disabling and sometimes fatal, but *P. leucopus* is remarkably resilient in the face of persistent infections with these pathogens, singly or in combination. How this species tolerates infections to otherwise thrive as well as it does is poorly understood.

*P. leucopus*’ importance as a pathogen reservoir, its resilience in the face of infection, and its appealing features as an animal model (6, 16), prompted our genetic characterization of this species, beginning with sequencing and annotating its nuclear and mitochondrial genomes (17, 18). The present study represents the third leg of this project, namely the microbial portion of the total animal “genome” for this species. Given the development of bait-delivered oral vaccines targeting *P. leucopus* (19) and plans to genetically modify and release this species (20, 21), pushing ahead on these interventional fronts without better understanding *Peromyscus* microbiota, the gastrointestinal (GI) tract’s in particular, seemed shortsighted.

Accordingly, we carried out a combined microbiologic and metagenomic study of the GI microbiome of *P. leucopus*. Our study focused on animals of a stock colony that has for many years been the major source of animals for different laboratories and spin-off breeding programs, including our own, in North America. The study extended to samples of *P. leucopus* deermice in their natural environments and, for a comparative animal, vivarium-reared *Mus musculus* under similar husbandry. While our investigations revealed similarities between the microbiota of the white-footed deermouse and the house mouse, there were also notable differences. These included a greater abundance and diversity of lactobacilli in *P. leucopus*. The investigation of four *Lactobacillus* species, particularly in their niches in the stomach of *P. leucopus*, was a special emphasis. A comparison of the GI microbiota of a natural population of *P. leucopus* and the stock colony animals revealed several species in common, albeit with larger variance among the wild animals.

## Results and discussion

### High coverage sequencing of fecal metagenome

Since there was only limited information in the literature on the GI microbiota of *P. leucopus* (22, 23), we began with untargeted assessment of microbiome constituents and their diversity in a sample of fecal pellets collected from two adult males and two adult females of the same birth cohort and shipment.

DNA was extracted and used for library construction; 332,279,332 paired-end reads of average length 247 nt were obtained after quality control and trimming of adapters. The mean % GC content was 47; 90% of the trimmed reads had PHRED scores of ≥ 30. The reads were characterized as to families of bacteria, parasites, and DNA viruses at the metagenomic server MG-RAST (http://mgrast.org). Annotated proteins accounted for 65% of the reads, followed by unknown proteins at 34%, and then ribosomal RNA (rRNA) genes at 0.8%. The rarefaction curve became asymptotic at 200,000 reads and a species count of 9000 (Fig S1 of Supplementary information). The alpha diversity was 250 species. By phylum 94% of the matched reads were either Bacteroidetes (60%) or Firmicutes (34%) (Fig S2 of Supplementary information). Higher level functional categories included carbohydrates (16.4% of reads), clustering-based subsystems (14.8%), protein metabolism (8.9%), amino acids and derivatives (7.8%), RNA metabolism (6.6%), and DNA metabolism (5.7%) (Fig S3 of Supplementary information).

A portion of the DNA was also submitted for commercial 16S rRNA metagenomics analysis of microbiota. As illustrated in Fig 1 and detailed in Tables S1 and S2 of Supplementary information, for the 20 most abundant taxa at the family or higher, there was concordance between the methods in the rankings. The most common families by the metagenomic accounting were members of the gram-negative bacterial order Bacterioidales (*Bacterioidaceae*, *Porphyromonadaceae*, *Prevotellaceae*, and *Rikenellaceae*), the gram-positive phylum Actinobacteria (*Bifidobacteriaceae* and *Coriobacteriaceae*), or the gram-positive phylum Firmicutes (*Bacillaceae*, *Enterococcaceae*, *Lactobacillaceae*, unclassified Clostridiales, *Eubacteriaceae*, *Lachnospiraceae*, *Peptococcaceae*, *Ruminococcoceae*, Thermoanaerobacterales Family III, *Erysipelotrichaceae*, and *Veillonellaceae*). Two exceptions were organisms of the families *Spirochaetaceae* of the phylum Spirochaetes and *Helicobacteraceae* of the phylum Proteobacteria.

**Fig 1.**
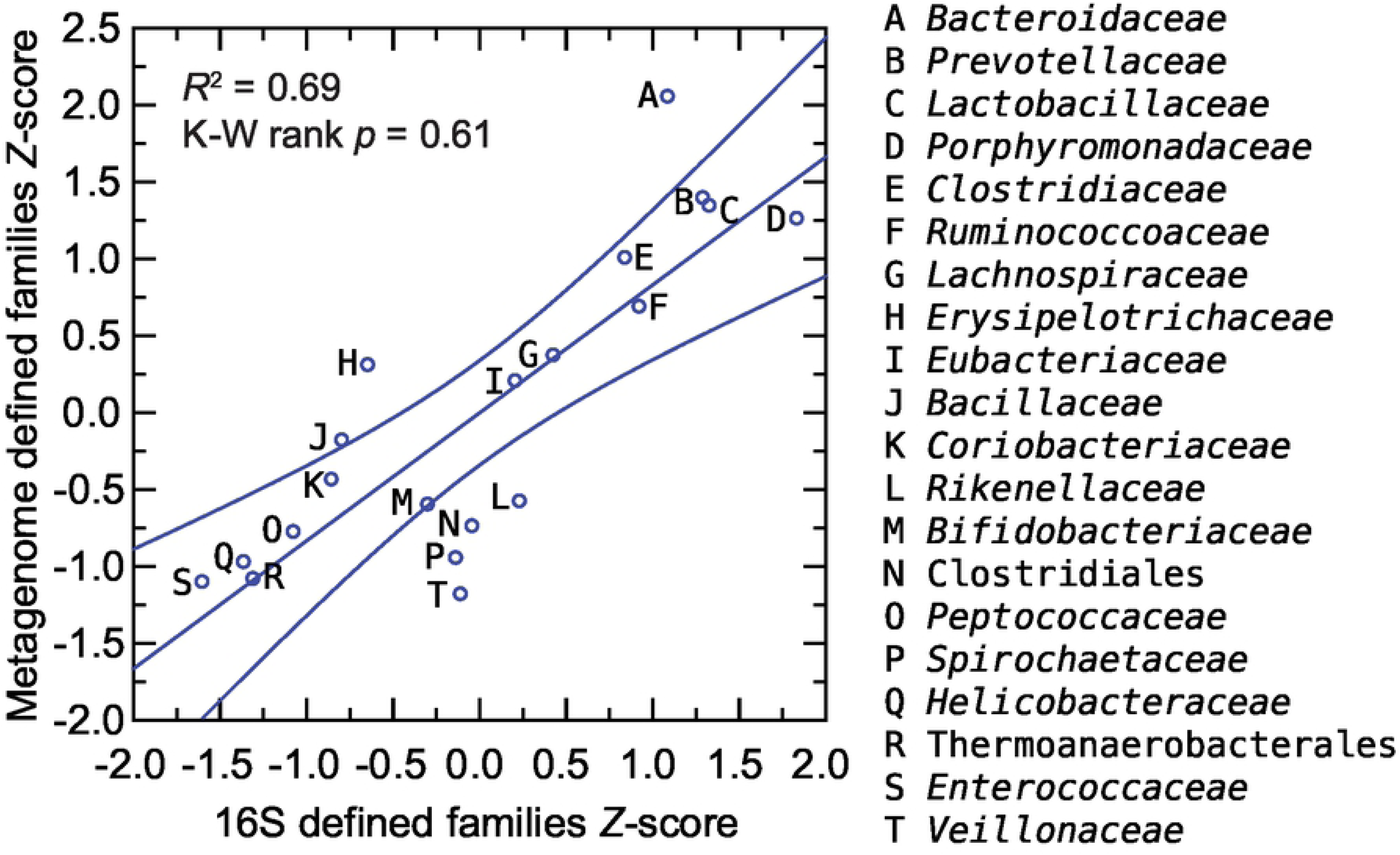
Scatter plot of relative abundances of commonly occurring bacterial families or orders in fecal metagenomes of Peromyscus leucopus LL stock by 16S ribosomal RNA gene criteria (x-axis) and by genome-wide gene (y-axis). The values of the two methods were normalized by Z-score. The different taxa are indicated in the graph capital letters, which defined in the box to right. The linear regression curve and its 95% confidence interval is shown. The coefficient of determination (R2) value and the Kruskal-Wallis (K-W) test by ranks p value are given.

A de novo assembly yielded 16,945 ungapped contigs of ≥ 10 kb from 197,369,943 reads and totaling 385 Mb of sequence with an average coverage of 104X. Of the total, 219 contigs were ≥ 100 kb in length and with ≥ 30X coverage. These were used in searches of non-redundant nucleotide and protein databases for provisional classifications. The identified taxa included Bacteroidales, Clostridia, *Clostridiaceae*, Erysipelotrichales, *Lactobacillaceae*, *Muribaculaceae*, Firmicutes, and *Spirochaetaceae*. Three organisms represented among the high coverage contigs could be unambiguously classified as to species: *Lactobacillus animalis*, which is considered in detail below, and two *Parabacteroides* species: *distasonis* and *johnsonii*. Another *Lactobacillus* species represented among the highly ranked contigs could not be identified with a known species represented in the database (24).

Among the high coverage contigs were also representatives of Rhodospirillales of the class Alphaproteobacteria, *Mycoplasmataceae* of Mollicutes, and the little-characterized group of bacteria called Elusimicrobia (25). Nearly as prevalent were organisms closely related to the phylum-level designation *Candidatus* Melainabacteria (26). On the list of organisms identified by searches with metagenomic contigs of databases and cumulatively accounting for 95% of the matched reads were the unexpected finding of the protozoan taxon *Trichomonadidae* with 41,614 or 0.12% of the reads (Table S2 of Supplementary information). *Enterobacteriaceae* at 0.3% accounted for a relatively small proportion of matched reads.

As in humans (27), *Bacteroidaceae*, *Lachnospiraceae*, Prevotellaceae, and *Ruminococcaceae* were abundant in the gut metagenome and cumulatively accounted for approximately half of the identified families in the *P. leucopu*s sample. One difference between humans and this *P. leucopus* sample was the much higher prevalence in *P. leucopus* of the family *Lactobacillaceae*, which on average represented only ∼0.2% of the metagenome in a European population (27) and by 16S sequencing ≤0.4% on the fecal microbiota in other studies (28). A higher proportion of lactobacilli in the fecal microbiota was previously noted in other rodents (29).

### Selected taxa

#### Escherichia coli

Although *Enterobacteriaceae* were infrequently represented among the metagenomic sequences, their cultivability under routine laboratory conditions and the availability of a vast database prompted our isolation of *Enterobacteriaceae* from LL stock *P. leucopus* fecal pellets on selective media. The predominant isolate on the plates was an *Escherichia coli*, which we designated LL2. The whole genome sequence of isolate LL2’s chromosome and plasmids was sequenced and assembled using a hybrid of long reads and short reads (Table 1) for an overall coverage of 90X. The chromosome in two contigs of 3.4 Mb and 1.6 Mb totaled 5.0 Mb with a GC content of 50%.

**Table 1.**
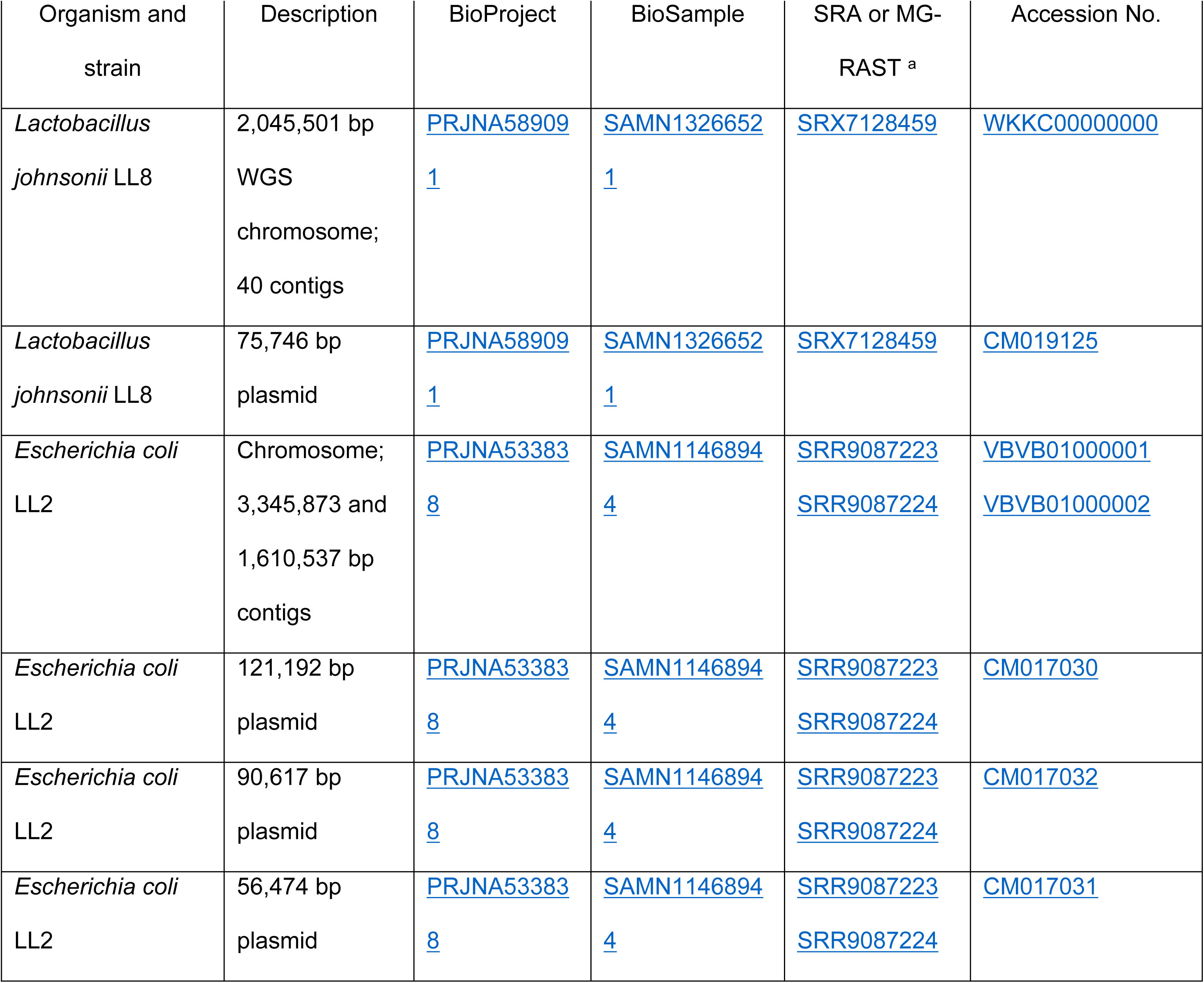
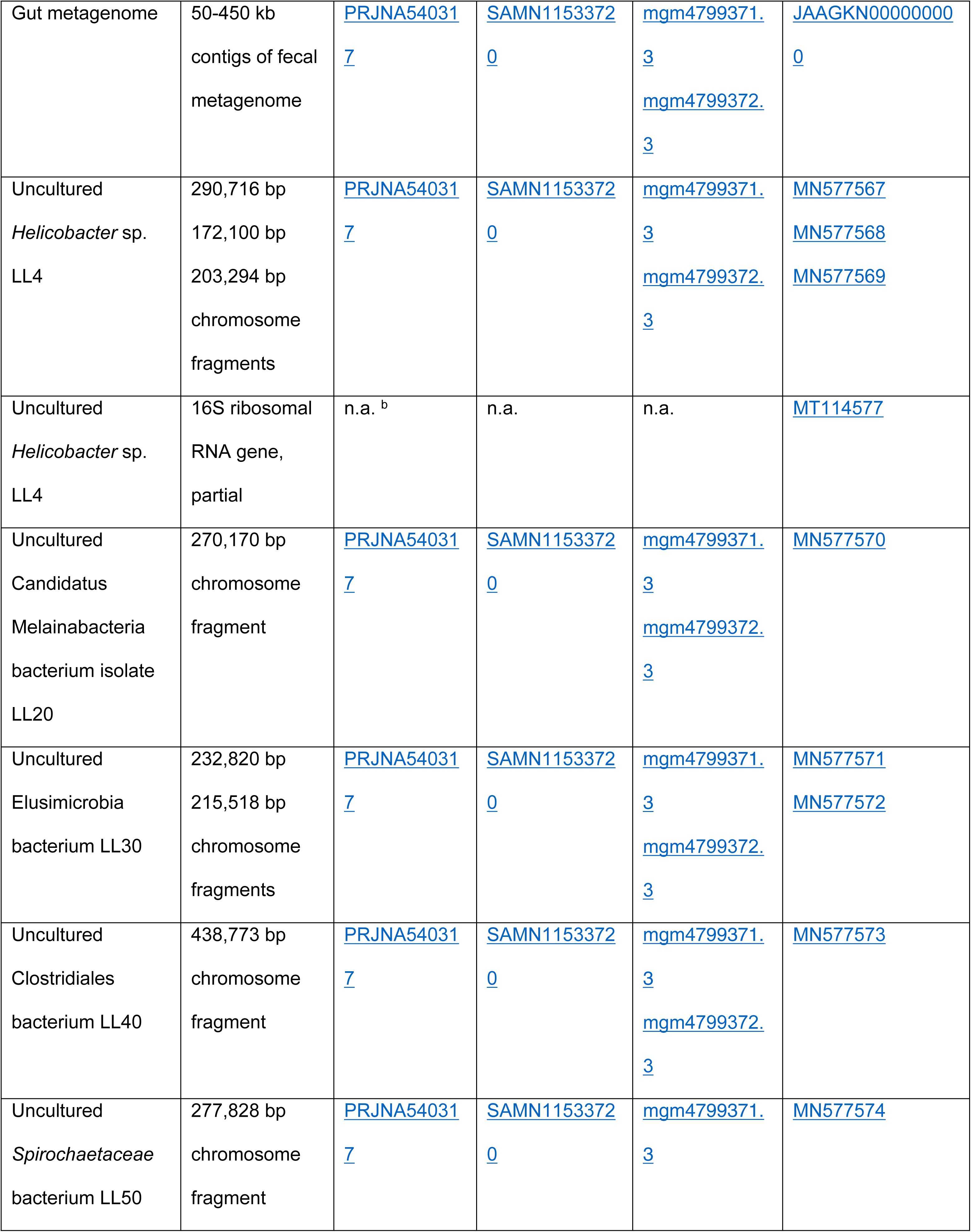
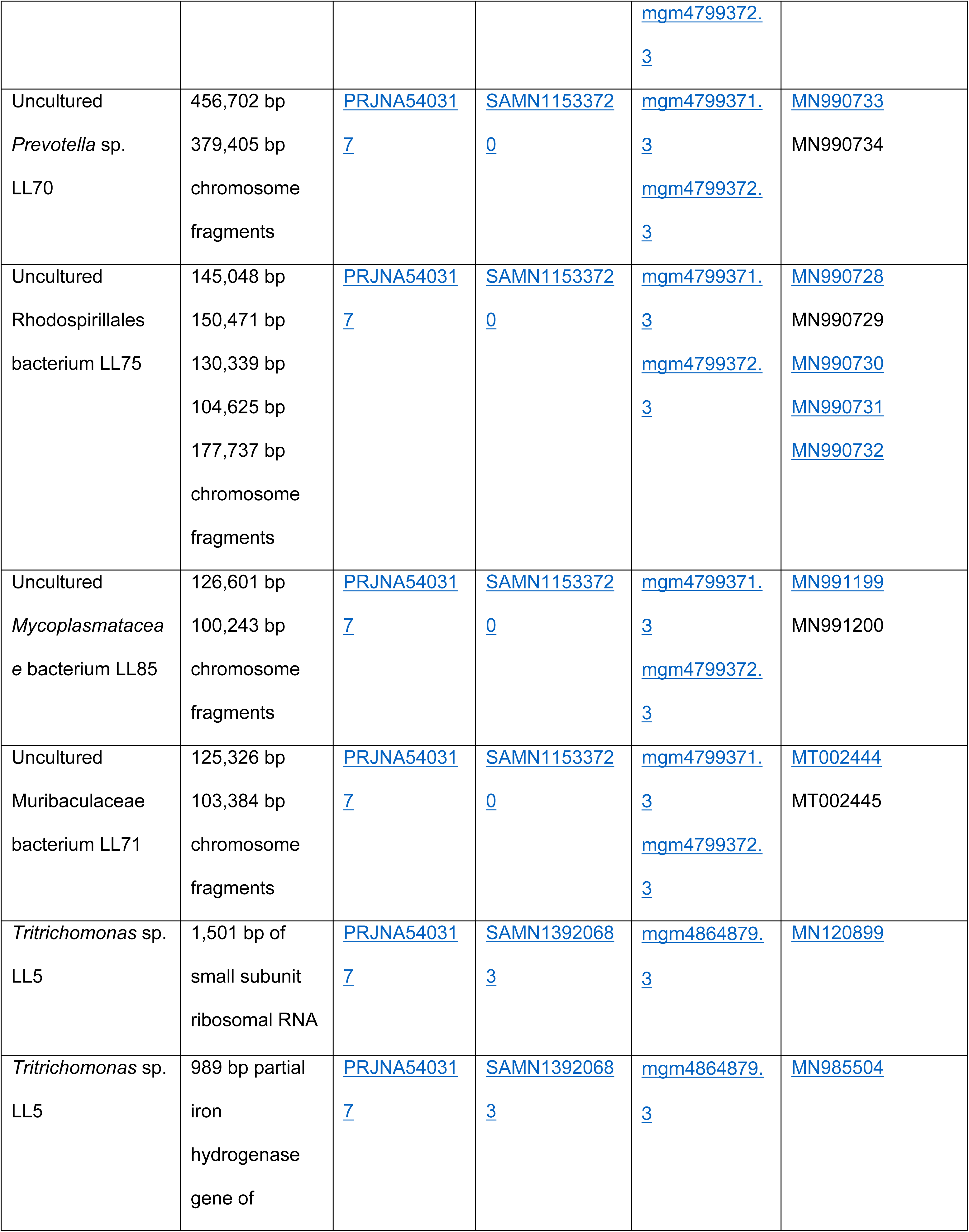
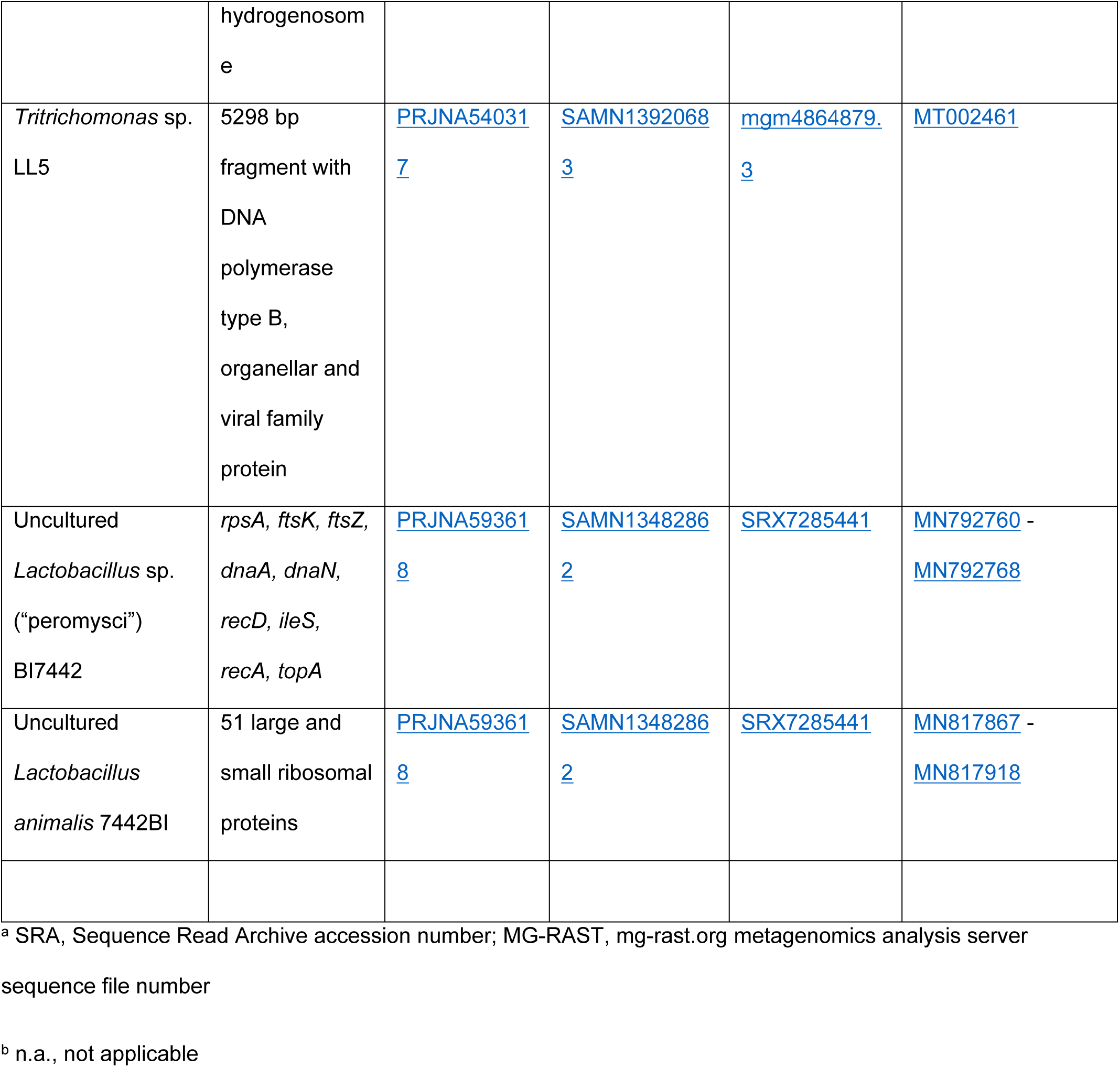
Resources from this study

*E. coli* LL2 had the following MLST schema types (http://pubmlst.org or http://enterobase.org): Achtman ST-278, Pasteur ST-357, and ribosomal protein ST-122394. The ribosomal protein profile was unique among thousands of isolates in the database. The 121 kb, 56.5 kb, and 91 kb plasmids of strain LL2 were similar to the following *E coli* plasmids, respectively: a 185 kb plasmid (NC_007675) found in an avian strain, a 58 kb plasmid (CP024858) of a multiply antibiotic-resistant human isolate, and an 89 kb plasmid (CM007643) in an organism isolated from sewage. *E. coli* LL2 was susceptible to ampicillin, ciprofloxacin, gentamicin, and sulfamethoxazole-trimethoprim by in vitro testing.

The chromosome was notable for the following: CRISPR-Cas1 and –Cas3 arrays; ISas1, ISNCY, IS3, IS110 and IS200 family transposases; restriction-modification systems; fimbria and curli biosynthesis and transport systems; type II toxin-antitoxin systems; and type II, type III and type VI secretion systems. The plasmids encoded fimbrial and pilin proteins, type I, type II, and type IV secretion systems, colicins, CdiA-type contact-dependent inhibition toxin, and three conjugative transfer systems, but no discernible coding sequences for antibiotic resistance.

Serial dilutions of feces of LL stock 20 animals (11 females and 9 males) in phosphate-buffered saline and plated on agar selective for gram-negative enteric bacteria yielded a mean (asymmetric 95% confidence interval) of 3,491 (677-18,010) colony-forming units (cfu) of *E. coli* per g of feces. This low density was consistent with the findings from metagenomic sequencing.

While the origin of this *E. coli* strain in the colony animals is obscure, it appears to be stably maintained among the gut microbiota of this population of *P. leucopus*. This adaptation may make it a candidate as a vector for delivering oral vaccines to this species (30).

#### Lactobacillus

We isolated lactobacilli from fecal pellets of stock colony *P. leucopus* on plates of selective medium that were incubated under microaerophilic and hypercapnic conditions at 37 °C. Four different species were identified. The genomes of three of organisms, namely *L. animalis* strain LL1, *L. reuteri* strain LL7, and a new species, designated as *Lactobacillus* sp. LL6 and provisionally named as “L. peromysci”, have been reported (24). The fourth genome, of the LL8 strain of *L. johnsonii*, is described first here (Table 1). *L. johnsonii*’s chromosome from cumulative contigs was 2,045,501 bp, about the same size as that of “L. peromysci” at 2,067,236 bp, but shorter than the 2,280,577 bp length for *L. animalis* and 2,205,740 bp for *L. reuteri*. The % GC content of “L. peromysci” at 33.5 was closer to *L. johnsonii* (34.4) than to either *L. animalis* (41.0) or *L. reuteri* (38.9).

Fig 2 is a distance phylogram of 1385 aligned sites of 16S ribosomal RNA genes for the four different lactobacilli. These were distributed across four major groups of the genus *Lactobacillus*. The phylogenetic relationships were examined in more depth by multilocus sequence typing of the 53 genes for ribosomal proteins. These were identified in the genomes, compared with other deposited sequences in the ribosomal MLST database (https://pubmlst.org) (31), concatenated, and then aligned with analogously concatenated DNA sequences from related species (Table S3 of Supplementary information). Bacteria with identical sequences for the 53 ribosomal protein genes were not found in the rMLST database of 133,460 profiles. The % GC contents of the concatenated coding sequences were 39.5, 40.8, 42.2, and 42.3 for *L. johnsonii*, “L. peromysci”, *L reuteri*, and *L. animalis*, respectively. Fig 3 shows the distance phylograms for ∼20 kb of aligned positions for the four species, each grouped with other strains or species within their respective phylogenetic clusters.

**Fig 2.**
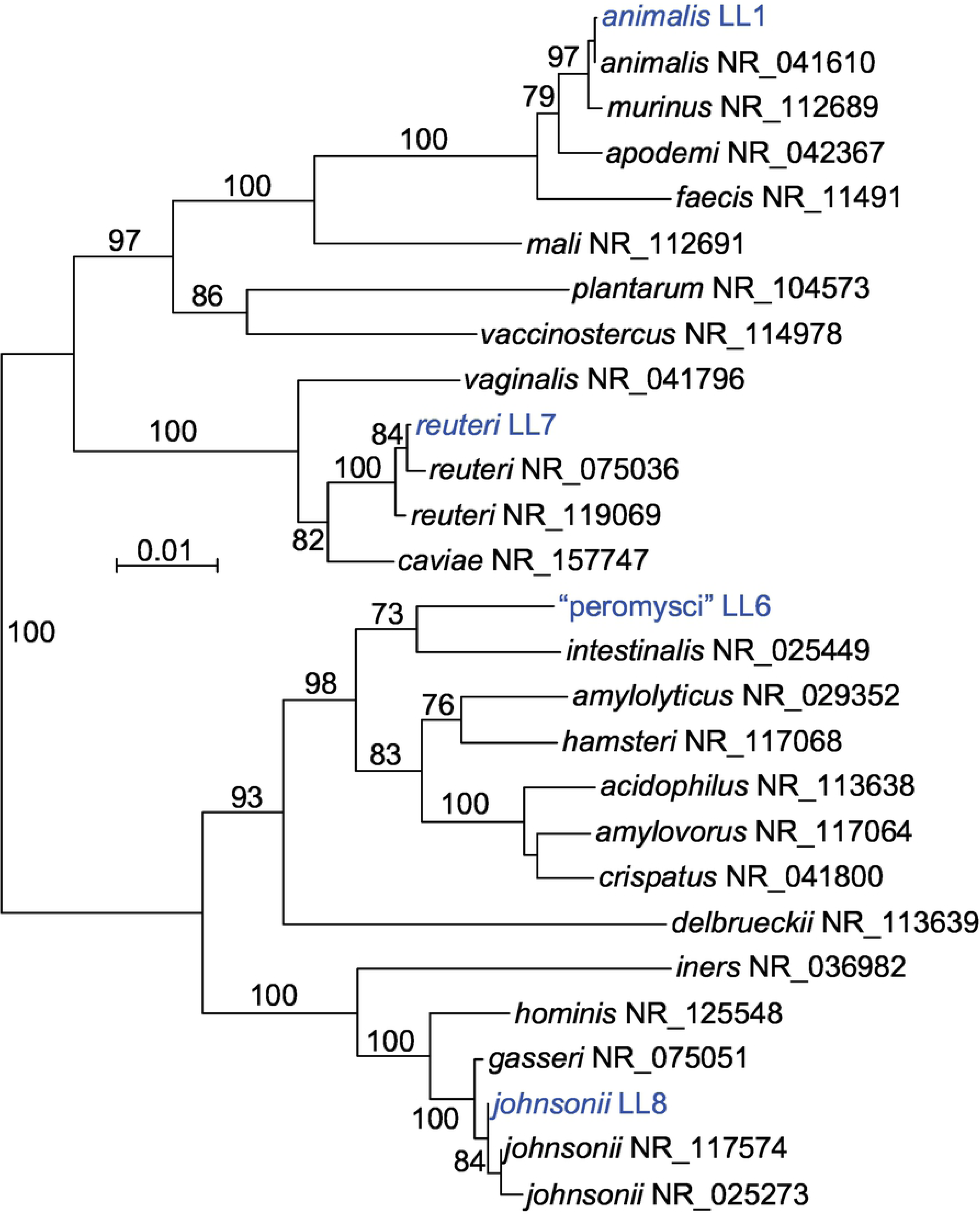
Neighbor-joining distance phylogram of 1420 aligned positions of 16S ribosomal RNA genes of the culture isolates of four *Lactobacillus* species from *Peromyscus leucopus* and selected other *Lactobacillus* spp. The sources for the accession numbers for the strains are given in Methods (*L. animalis*, *L. reuteri*, and “L. peromysci”) or in Table 1. The other organisms represented are from Reference RNA sequences database of the National Center for Biotechnology Information; the accession numbers are given after the species name. The scale for distance by criterion of observed differences is indicated. Percent bootstrap (100 iterations) support values of ≥ 90% at a node are shown.

**Fig 3.**
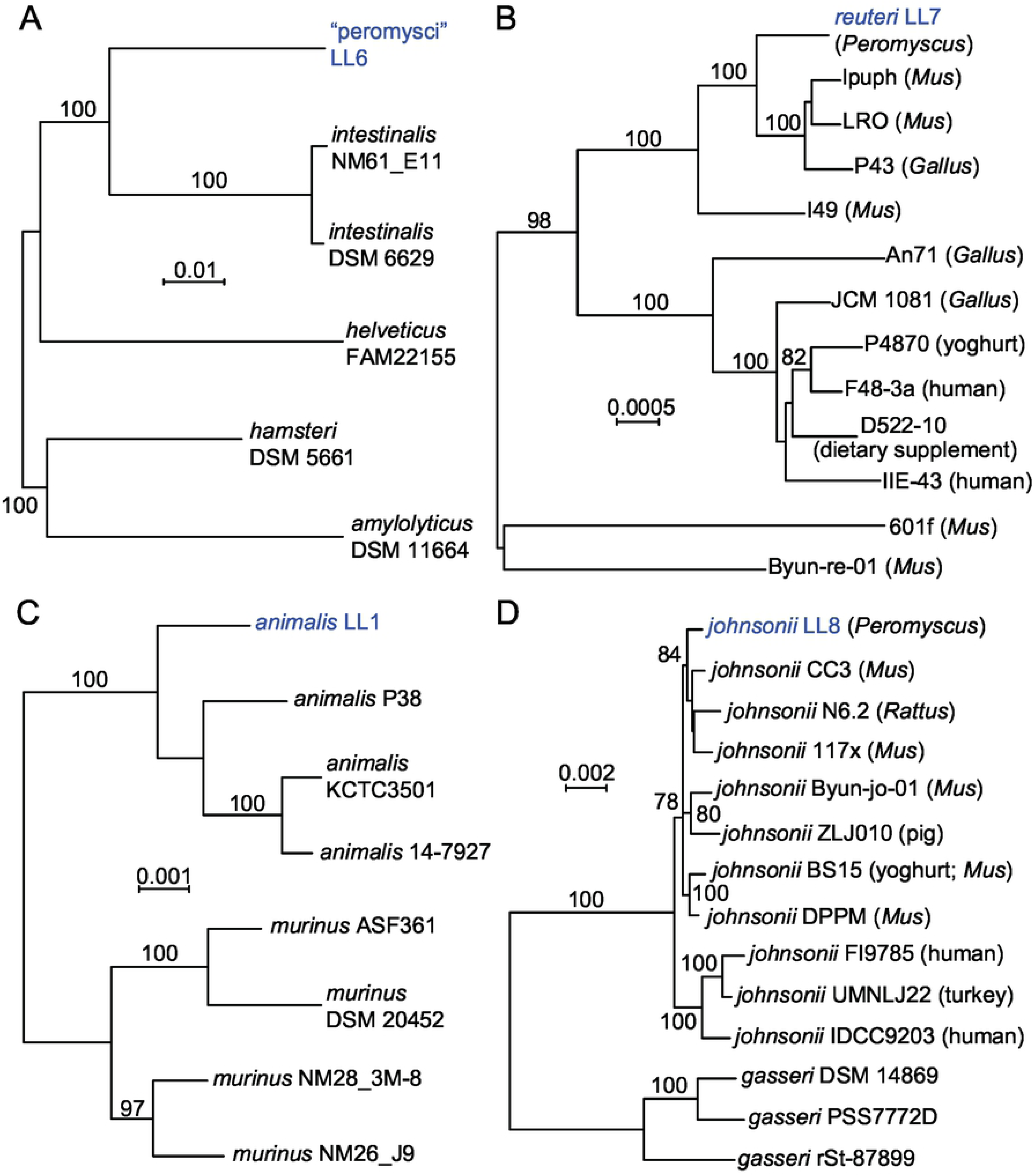
Neighbor-joining distance phylograms of codon-aligned, concatenated nucleotide sequences for complete sets of ribosomal proteins of “L. peromysci” (panel A), *L. reuteri* (panel B), *L. animalis* (panel C), and *L. johnsonii* (panel D) of *P. leucopus* compared with *Lactobacillus* spp. (strain identifier) of other sources. The scales for distance by Jukes-Cantor criterion are indicated in each panel. Percent bootstrap (100 iterations) support values of ≥ 75% at a node are shown. In panels B and D the host animal or other origin for a given isolate are given in parentheses.

“Lactobacillus peromysci” was distant from other sequenced lactobacilli by rMLST (panel A), as well as by its 16S ribosomal RNA gene (Fig 2). The nearest taxon in the sequence alignment was *L. intestinalis*, which was first isolated from the intestines of *Mus musculus* and other murids (32). The unique ST for the rMLST for strain LL6 of this organism is 115326.

Draft and complete genomes of numerous *L. reuteri* strains have been sequenced, for example, strain Byun-re-01, which was isolated from *M. musculus* small intestine (33). Many of these are utilized in the fermented foods industry, such as production of kimchi, or as dietary supplements, but others were isolated as constituents of the GI microbiota of several varieties of animals. *L. reuteri* strain LL7 was in a cluster that mainly comprised isolates from *M. musculus* (panel B).

*L. animalis* and *L. murinus* are closely related species that primarily have been associated with GI microbiota of rodents and some other mammals. Isolate LL1 grouped with representatives of *L. animalis* in the analysis (panel C) and not *L. murinus* (34). LL1’s 16 ribosomal RNA sequence was identical to that of the type strain ATCC 35046 of *L. animalis* (35) at 1488 of 1489 positions (GCA_000183825) (36). Another pair of closely-related species are *L. johnsonii* and *L. gasseri*, for which there are several sequenced genomes. The LL8 isolate from fecal pellets of *P. leucopus* clustered with *L. johnsonii* strains from mice and rats (panel D). More distant were strains of *L. johnsonii* isolated from humans and a bird; more distant still were representatives of *L. gasseri*.

Plasmids were identified in each of the four species on the basis of a circularly permuted sequence for a contig and presence of coding sequences that were homologous to known plasmid replication or partition proteins (Table 1). Large plasmids of 179 kb and 76 kb were present in *L. reuteri* and *L. johnsonii*, respectively. *L. animalis* and “L. peromysci” had small plasmids of 4 kb and 7 kb, respectively. Megaplasmids of greater than 100 kb have been observed in other *Lactobacillus* spp. (37). In all genomes there was evidence of lysogenic bacteriophages or their remnants. All species except *L. reuteri* discernibly had coding sequences for Class I or Class III bacteriocins or their specific transport and immunity proteins (Table S4 of Supplementary information).

Table 2 summarizes differentiating genetic profiles among the four species for 11 selected genes or pathways. Two species, *L. reuteri* and “L. peromysci”, had coding sequences for a urease, which could provide for tolerance of acidic conditions, such as in the stomach. A urease had previously been identified in a *L. reuteri* strain that was considered a gut symbiont in rodents (38). The four species had *secY1-secA1* transport and secretion systems. Accessory Sec systems (*secY2-secA2*) were identified in genomes of *L. reuteri*, *L. johnsonii*, and *L. animalis* but not in “L. peromysci”. The LL7 strain of *L. reuteri* on its megaplasmid also had coding sequences for a third SecY-SecA system. An accessory Sec system was involved with adhesion and biofilm formation in the Lactobacillales bacterium *Streptococcus pneumoniae* (39). A coding sequence for an IgA protease was identified in *L. johnsonii* but not in the other three species. An IgA protease in another strain of *L. johnsonii* was associated with long-term persistence in the gut of mice (40). The presence or absence of other genes or pathways that differentiated between the four species were an L-rhamnose biosynthesis pathway in one species, a *luxS* gene associated with a quorum sensing system in *L. reuteri* and *L. johnsonii* (41), a type 1 CRISPR-Cas3 array in “L. peromysci” (42), pathways for thiamine biosynthesis (43) and for reduction of nitrate (44) in three species, an arginine deiminase and its repressor in *L. reuteri* (45), and a type VII secretion system in *L. animalis* (46).

**Table 2.** Selected genes and pathways in four species of *Lactobacillus* of the gastrointestinal microbiota of *Peromyscus leucopus*

|  |  |  |  |  |  |  |  |  |  |  |  |
| --- | --- | --- | --- | --- | --- | --- | --- | --- | --- | --- | --- |
| Species | Urea<br>se | secY<br>2-secA<br>2 | secY<br>3-secA<br>3 | IgA<br>protease<br>(pfam07<br>580) | L-rhamn<br>ose<br>pathwa<br>y | lux<br>S | Type<br>1<br>CRIS<br>PR<br>Cas | Thiamin<br>e<br>biosynth<br>esis | Nitroreduc<br>tase<br>*nfnB-nifU* | Arginin<br>e<br>deimina<br>se/<br>repress<br>or | Type<br>VII<br>secreti<br>on<br>syste<br>m |
| *reuteri* | + | + | + | - | - | + | - | - | - | + | - |
| *johnsonii* | - | + | - | + | - | + | - | + | + | - | - |
| *animalis* | - | + | - | - | - | - | - | + | + | - | + |
| “peromysci” | + | - | - | - | + | - | + | + | + | - | - |

Of the four species found in *P. leucopus* feces, only *L. johnsonii* and *L. reuteri* have been commonly isolated from human feces (28). While various strains of *L. reuteri*, *L. johnsonii*, and either *L. animalis* or the closely related *L. murinus* have been observed among the GI microbiota of *M. musculus* represenstatives (47), an organism similar to “L. peromysci” has not. Whether this is an indication of a restricted host range or a specific adaptation for this bacterium is considered below.

#### Helicobacter

Among the assembled metagenomic contigs were three totaling 666,100 bp of a *Helicobacter* genome (Table 1). The contigs had non-overlapping in genetic content, and blast searches with translated genes from each of the 3 contigs yielded the identical rankings of taxa for homologous proteins. On these bases, we concluded that the contigs represented a single type of *Helicobacter* bacterium, and designated it strain LL4. Using the DNA sequences for 53 ribosomal proteins of this organism, we compared it with similar sets from other *Helicobacter* spp. (panel A of Fig 4;). This analysis, as well as analysis of the 16S ribosomal RNA gene sequence from a fecal sample from another animal (Fig S5 of Supplementary information), showed that the organism was near-identical to orthologous sequences of *Helicobacter* sp. MIT 05-5293 (accession JROZ02000000), which had been cultivated from a wild *P. leucopus* captured in Massachusetts (48; J.G. Fox, personal communication). This finding indicated that the organism was autochthonous for *Peromyscus* and had not been acquired from another rodent housed in the same vivarium. LL4 and MIT 05-5293 are in a cluster of species known as “enterohepatic” *Helicobacter* for their primary residence in the intestine rather than the stomach and for their frequent presence in liver tissue (49). These species may not be benign. *H. hepaticus* is associated with hepatitis, bowel inflammation, and carcinoma (50), and *H. typhlonius* is associated with reduced fecundity in mice (51).

**Fig 4.**
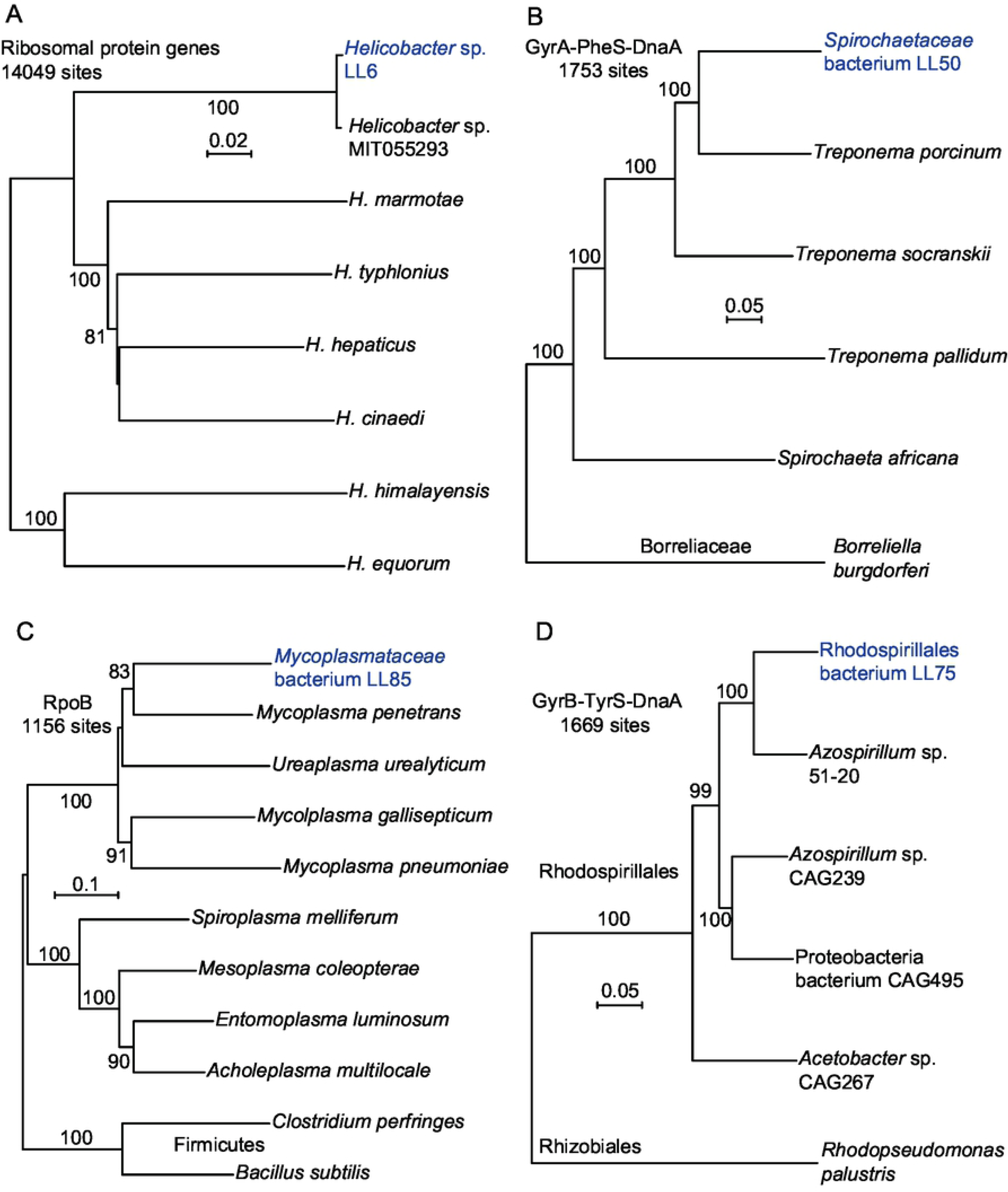
Neighbor-joining distance phylograms of concatenated nucleotide (panel A) or amino sequences (panels B-C) of Helicobacter spp. (panel A), *Spirochaetaceae* bacteria (panel B), Mollicutes and Firmicutes bacteria (panel C) and Rhodospirillales bacteria (panel D) of gut metagenome of *P. leucopus* and from other sources. The respective phylogenetic analyses used concatenated sequences of the following: ribosomal protein genes (panel A); the DNA gyrase A (GyrA), phenylalanyl t-RNA synthase, alpha subunit (PheS), and chromosomal replication iniator protein (DnaA) (panel B); DNA-directed RNA polymerase, beta-subunit (RpoB) (panel C); and DNA gyrase B (GyrB), tyrosyl t-RNA synthase (TyrS), and DnaA (panel D). The distance criteria were Jukes-Cantor for the codon-aligned nucleotide sequences and Poisson for amino acid sequences. The scales for distance are shown in each panel. Percent bootstrap (100 iterations) support values of ≥ 80% at a node are shown.

#### Spirochaetaceae

Panels B, C, and D of Fig 4 are phylograms of three other types of bacteria that were identified among the high-coverage metagenomic contigs (Tables 1 and S5). The uncultured spirochete LL50 was placed in the genus *Treponema* by the MG-RAST analysis program. Yet species in this genus are highly divergent and include free-living organisms in a variety of environments, symbionts of termites, the agent of syphilis, and gut residents, such as *T. porcinum*, which was isolated from the feces of pigs (52). More distant still was the agent of Lyme disease, *Borreliella burgdorferi*, of the family *Borreliaceae* (53). In our view naming the organism as a “treponeme” would provide little insight about its role in the microbiome and may even be misleading.

### Seven other bacterial taxa

The *Mycoplasmataceae* bacterium LL85 (panel C of Fig 4 and Table S5) was unlike any other mollicute represented in the database but was in cluster with vertebrate-associated species, like *M. pneumoniae*. But there is also deep branching in this tree, as the tree including as outgroup two Firmicutes shows. Panel D is a phylogram of selected alphaproteobacteria and includes the organism LL75 (Table S5). The algorithmic analysis identified this at the genus level as *Azospirillum*, which is a largely uncharacterized taxon with highly divergent members. While assignments as to genus or family are uncertain at this time, LL75 clustered within the order Rhodospirillales and not with rhizobacteria.

Table 1 lists five other types of novel bacteria that were identified in the *P. leucopus* gut metagenome and partially sequenced and annotated. These were a *Candidatus* Melainabacteria bacterium (isolate LL20), an Elusimicrobia bacteria (isolate LL30), a Clostridiales bacterium (isolate LL40), a *Prevotella* species (isolate LL70), and a *Muribaculaceae* bacterium (isolate LL71). *Candidatus* Melainabacteria is either a non-photosynthetic sister phylum of cyanobacteria or a class within the phylum Cyanobacteria (26). Besides a variety of environmental sources, including hot springs and microbial mats, these poorly-characterized organisms have also been identified in the feces of humans and other animals. The phylum Elusimicrobia, formerly “Termite Group 1” (54), is a strictly-anaerobic, deeply-branched lineage of gram-negative bacteria, representatives of which were first observed in the hindgut of termites (25). The family *Muribaculaceae* (formerly “family S24-7”) of the order Bacteroidales were first identified among gut microbiota of mice and subsequently in the intestines of other animals, including humans and ruminants (55).

### DNA viruses

Of 112,677,080 reads of the metagenome high-coverage sequencing of the LL stock animals, 97,147 (0.09%) were assigned to one of 28 DNA virus families. Three classifications accounted 92% of the reads: *Siphoviridae* (50%), which are bacteriophages with long contractile tails; *Myoviridae* (21%), which are bacteriophages with contractile tails; and “unclassified viruses” (21%). At the species level, 31,812 (68%) of the 46,904 *Siphoviridae*-matching reads mapped specifically to bacteriophages of *Lactobacillus* spp.

#### *Tritrichomonas* protozoan

Intestinal flagellated protozoa named “Trichomonas muris” or “Tritrichomonas muris” had previously been identified in wild *P. leucopus* and *P. maniculatus* (56). While laboratory mice are typically free of intestinal protozoa (57), the anaerobic *Tritrichomonas muris* has been reported in some populations of colony *M. musculus* (58). To further investigate the protozoa that were provisionally identified as “Trichomondidae” at the family level in the metagenome analysis, we euthanized 14 healthy adult animals (6 females and 8 males) and examined fresh cecal contents by phase microscopy. Six of the LL stock animals had been born at the PGSC facility, and 8 had been born at U.C. Irvine.

In each of the 14 animals examined there were numerous motile flagellates consistent in morphology with *T. muris* in their ceca (59). These were each at a cell density of ∼10^6^ per milliliter of unconcentrated cecal fluid (Fig 5; S1 File). Entire ceca and their contents from two adult females and two adult males were subjected to DNA extraction, library preparation from the DNA, sequencing, and de novo assembly of contigs.

**Fig 5.**
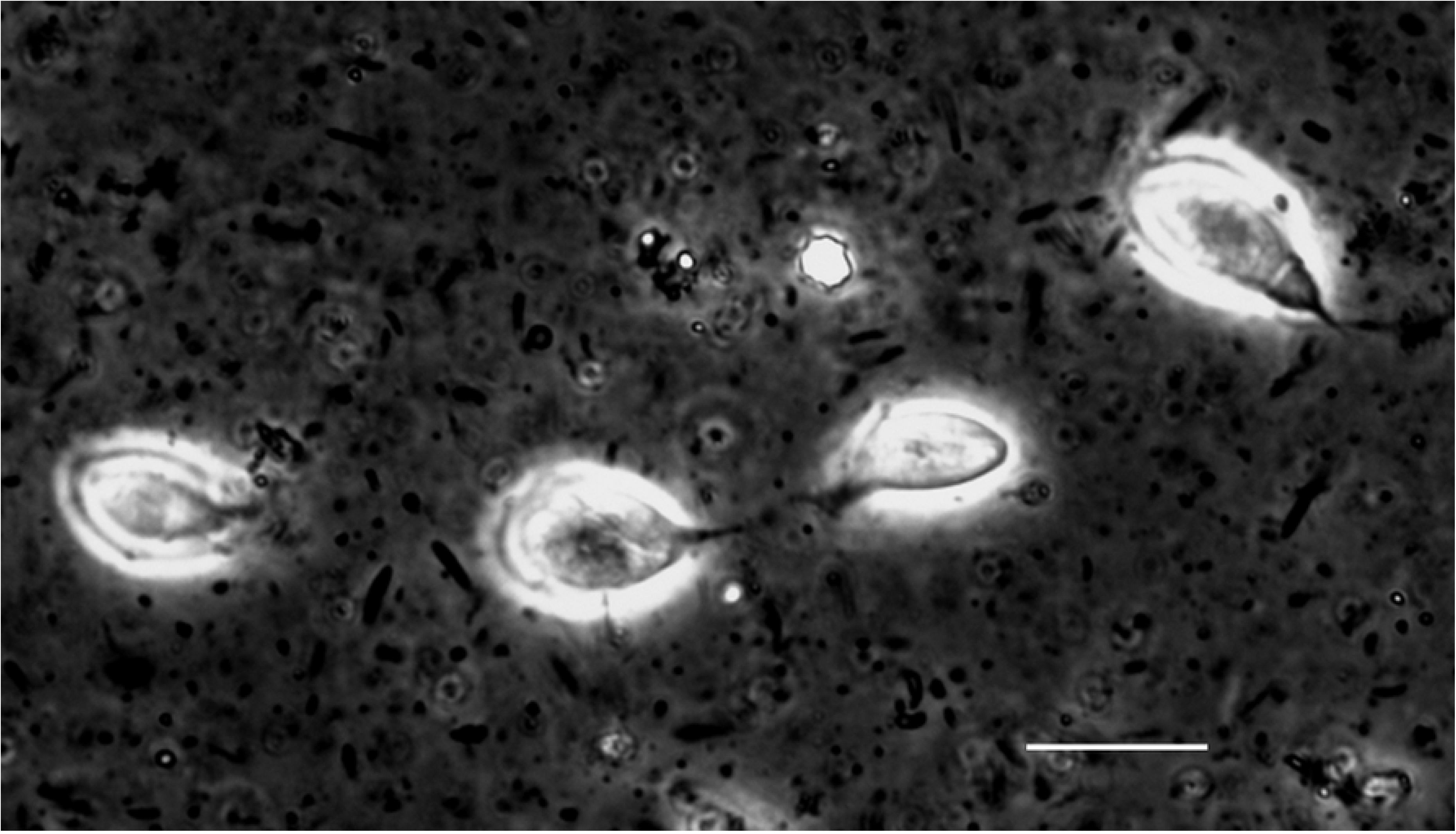
Photomicrograph of live *Tritrichomonas* flagellated protozoan in the cecal fluid of *P. leucopus* LL stock. Four organisms against the background of intestinal bacteria were visualized in the wet mount by differential interference microscopy. Bar, 10 µm.

Fig 6 shows phylograms of nucleotide sequence of the small subunit (SSU) ribosomal RNA gene (panel A; Table S5) and of the partial amino acid sequence of the iron hydrogenase of the hydrogenosome of anaerobic protozoa (panel B; Table S5) (60). The SSU of isolate LL5 indicates that it is probably synonymous with *Tritrichomonas muris*, for which only a SSU sequence was available. The sequence of the iron hydrogenase further supported placement in the genus *Tritrichomonas*. *Histomonas melagridis*, a sister taxon by this analysis, is recognized as a pathogen of poultry. Another sequence of the LL5 organism encodes a type B DNA polymerase (Table 1), which likewise matched closely with an ortholog in the *Tritrichomonas foetus* genome sequence (Fig S4 of Supplementary information). *T. foetus* is a sexually-transmitted pathogen of cattle (61) and a cause of chronic diarrhea in domestic cats (62).

**Fig 6.**
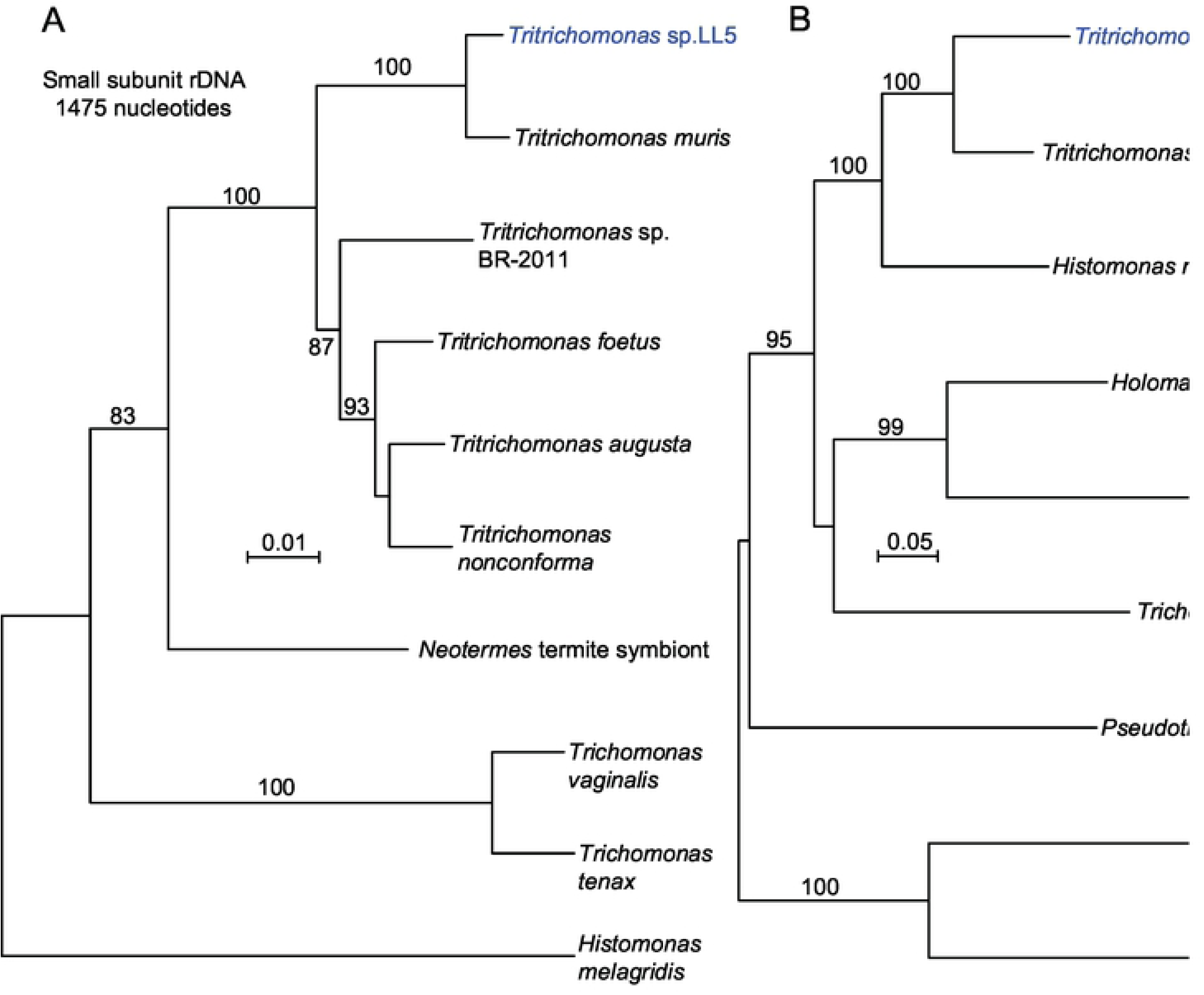
Neighbor-joining distance phylograms of nucleotide (panel A) or amino acid (panel B) partial sequences of small subunit ribosomal RNA gene (rDNA) (panel A) and iron hydrogenase protein (panel B) of *Tritrichomonas* sp. LL5 of *P. leucopus* and selected other parabasilids and other microbes. The distance criteria were observed differences for nucleotide alignment and Poisson for amino acid alignment. The scales for distance are shown in each panel. Percent bootstrap (100 iterations) support values of ≥ 80% at a node are shown.

Whether the *T. muris* is a commensal shared across natural populations of *Peromyscus* or a parasite acquired from another rodent during the colony’s history in a vivarium remains to be determined. As related below, there is sequence evidence of the same or related organism in several wild animals. Whatever the case, these organisms may have an effect on immune responses of *P. leucopus*, as has been reported for *T. muris* in *M. musculus* (63–65), and their presence needs to be taken into account in interpreting experimental results in the laboratory and in applications for field interventions.

### Comparative study of GI microbiota of *P. leucopus* and *M. musculus*

The preceding study revealed several microbes that were either undescribed species or genera, e.g. “L. peromysci” or the *Candidatus* Melainabacteria bacterium, or new strains of known microbial species, e.g. *L. animalis* LL1 and *T. muris* LL4. These novelties notwithstanding, to what extent did the gut microbiota of this deermouse resemble that of the typical laboratory animal, a house mouse that was maintained under similar husbandry conditions, including diet? That question motivated the following experiment.

Fecal pellet samples from 20 adult *P. leucopus* (10 females and 10 males) and 20 adult BALB/c *M. musculus* (10 females and 10 males) were obtained and stored frozen at −80 °C until processing. All animals were approximately 10 weeks old. The animals were housed in the same vivarium facility, though in different rooms. The pellets were subjected to total DNA extractions, and paired-end Illumina sequencing with 250 cycles of indexed libraries were carried out. There were means (95% CI) of 3.4 (3.1-3.7) x 10^6^ post-quality control reads for *P. leucopus* samples and 3.4 (3.2-3.6) x 10^6^ for *M. musculus* samples (Tables S6 and S7 of Supplementary information).

The reproducibility between replicate library constructions from the same sample was assessed with quantitations of reads assigned by taxonomic family for specimens from seven *P. leucopus* among the 20 total. Pairwise coefficients of determination (*R*^2^) for the 91 possible combinations were calculated (Table S8 of Supplementary information). The mean (95% CI) of *R*^2^ values were 0.999 (0.999-1.0) for the 7 pairs of replicates and 0.930 (0.915-0.944) for the 84 non-replicate pairs. We concluded that most of the variation between samples was attributable to inter-specimen differences in the microbiota and not to technical issues in library preparation or sequencing.

The prevalences of different taxonomic families in the *P. leucopus* and *M. musculus* gut metagenomes were similar (left panel of Fig 7 and Table S9 of Supplementary information). But a few families stood out as either more or less common in the deermice. Notable among these were *Lactobacillaceae*, *Helicobacteriaceae*, and *Spirochaetaceae*, which were approximately 4x, 8x, and 2x, respectively, more prevalent on average among microbiota of *P. leucopus* than in *M. musculus*. There was no evidence of *Tritrichomonas* sp. in the BALB/c mice by this analysis, but direct examination of intestinal contents was not carried out.

**Fig 7.**
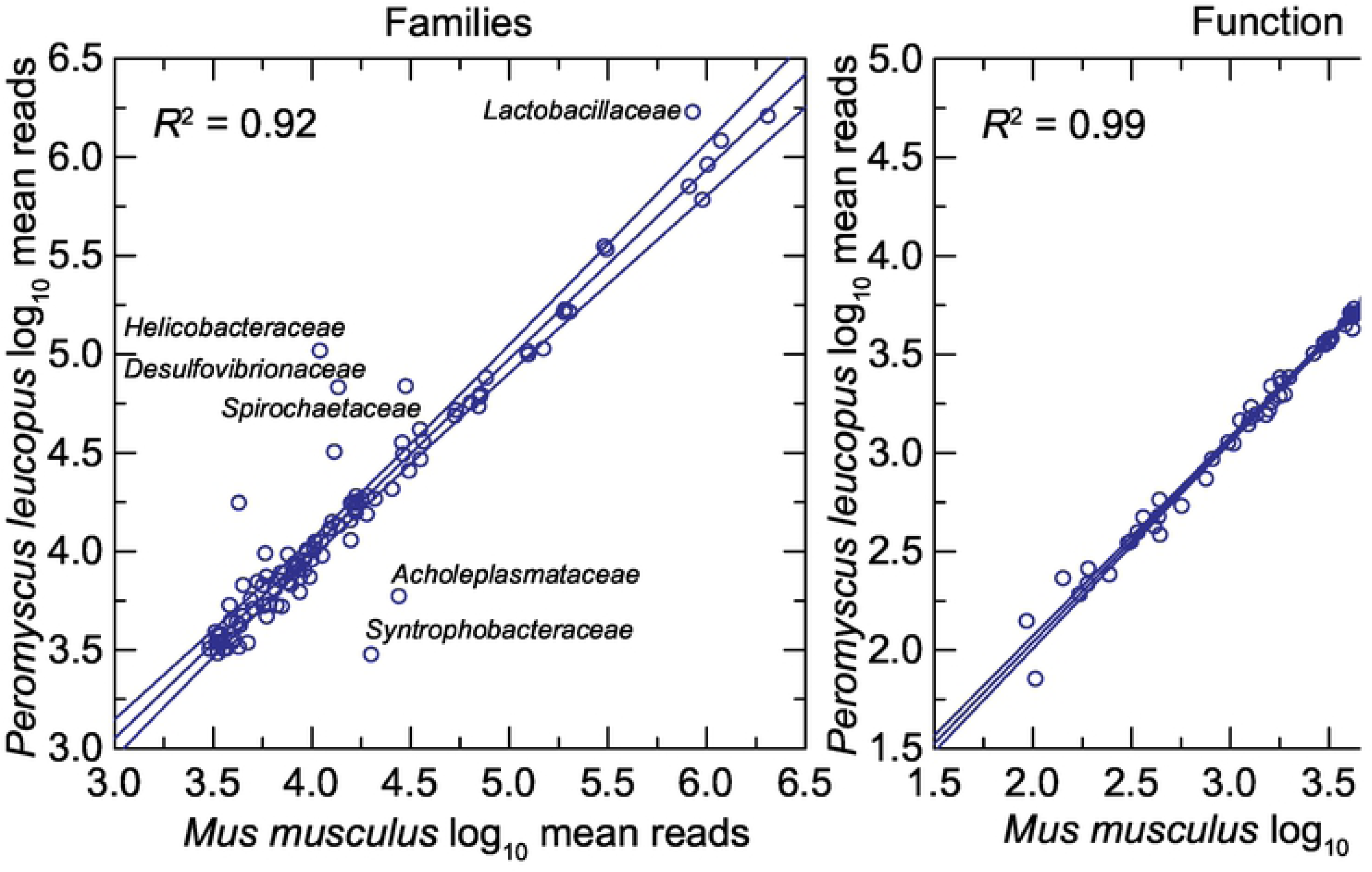
Scatter plots of log-transformed normalized reads of the gut metagenomes of 20 *P. leucopus* on the gut metagenomes of 20 *M. musculus* by bacterial families (left panel) or by function at the pathway level (right panel). The linear regression lines, their 95% confidence intervals, and coefficients of determination (*R*^2^) are shown. Selected families that are comparatively more or less prevalent in *P. leucopus* are indicated.

At the level of 86 operational KEGG pathways, the metagenomes of *P. leucopus* and *M. musculus* were nearly indistinguishable (right panel of Fig 7 and Table S10 of Supplementary information). But, as shown in the heat map of Fig 8, at the homologous gene level there were many differences between these two species and also between females and males within each species (Tables S12 and S13 of Supplementary information). Hierarchical clusters 2 and 4 of the analysis discriminated between mice and deermice regardless of sex, while clusters 1 and 3 signified marked differences by sex and less so by species.

**Fig 8.**
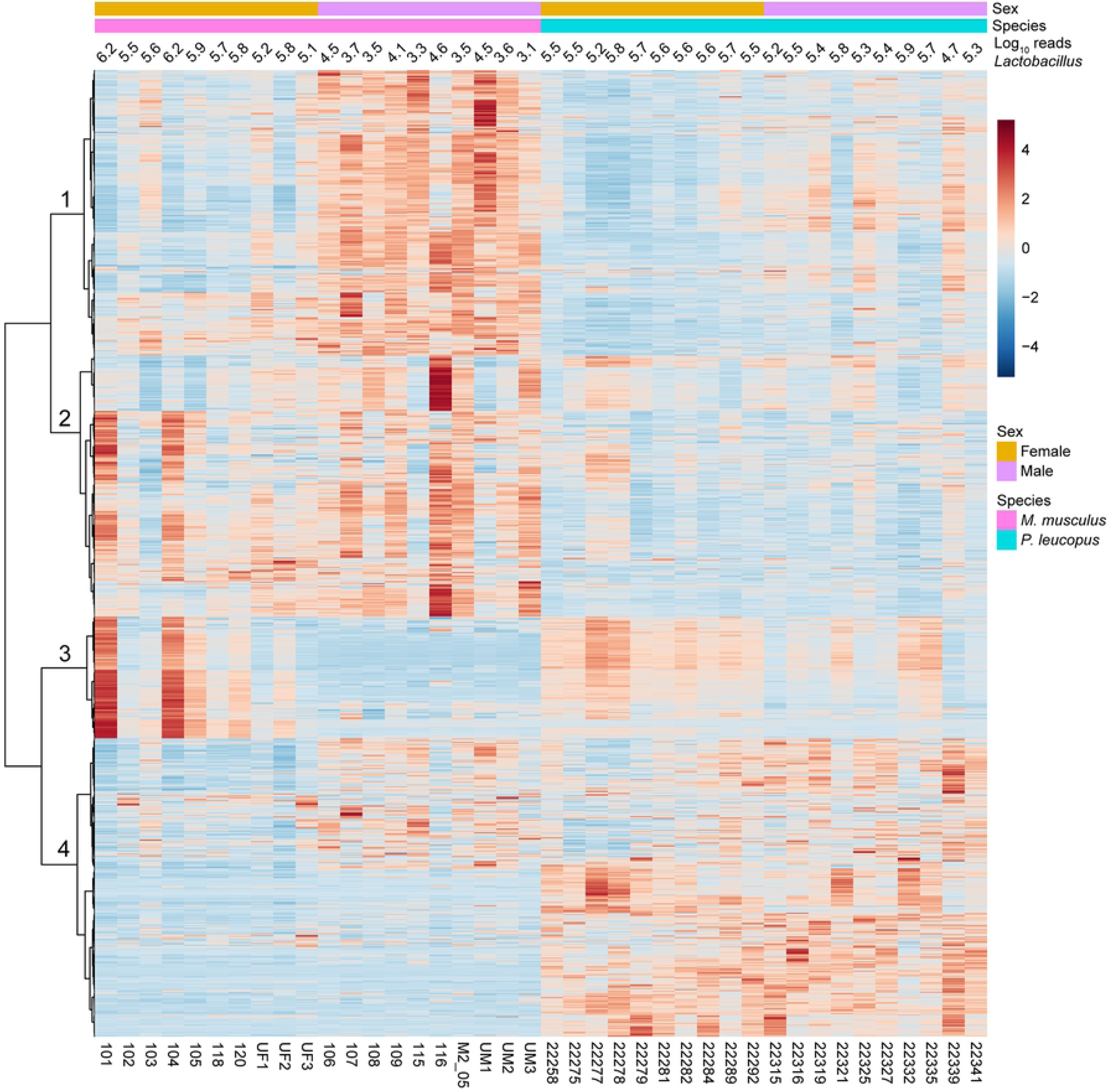
Heat map-formatted shading matrix of KEGG Orthology gene level annotations of gut metagenomes of *P. leucopus* and *M. musculus*. The annotations were generated by MicrobiomeAnalyst (https://www.microbiomeanalyst.ca). Columns are grouped by species and by sex within each species. Individual animal identifications are given on the *x*-axis below the heat map. Above the heat map are the log-transformed reads mapping to the genus Lactobacillus for each animal’s fecal sample. Clustering of rows of genes were by Pearson correlation coefficient. Four major clusters are labeled 1-4 on the *y*-axis. Scaling is by relative abundances from low (blue) to high (red).

As one example of differences between species, there was higher representation of genes of the mevalonate pathway in the gut metagenomes of *P. leucopus*. Beginning with acetyl-CoA and ending with isopentenyl pyrophosphate, the central intermediate in the biosynthesis of isoprenoids in all organisms (66), the coding sequences for the following ordered enzymes (with Enzyme Commission [EC] number) in the pathway were comparatively higher in frequency: acetyl-CoA C-acetyltransferase (EC:2.3.1.9), hydroxymethylglutaryl-CoA synthase (EC:2.3.3.10), hydroxymethylglutaryl-CoA reductase (EC:1.1.1.88), mevalonate kinase (EC:2.7.1.36), phosphomevalonate kinase (EC:2.7.4.2), and diphosphomevalonate decarboxylase (EC:4.1.1.33).

We further investigated specific differences between *P. leucopus* and *M. musculus* and between individual animals of each species in *Lactobacillus* spp. (67). This was achieved by mapping reads to references of the chromosome sequences of the four species that had been isolated from the feces of LL stock *P. leucopus*. The caveat is that the lactobacilli in the mice would not be expected to be identical to the deermouse strains used as references. Fig 9 shows box plots for *Peromyscus* on the left and for *Mus* on the right for data given in Table S11 of the Supplementary information. Included in the analysis of *P. leucopus* gut metagenome reads were selected other bacteria that had been frequently identified among the metagenomic contigs and then further characterized by partial genome sequencing (see above).

**Fig 9.**
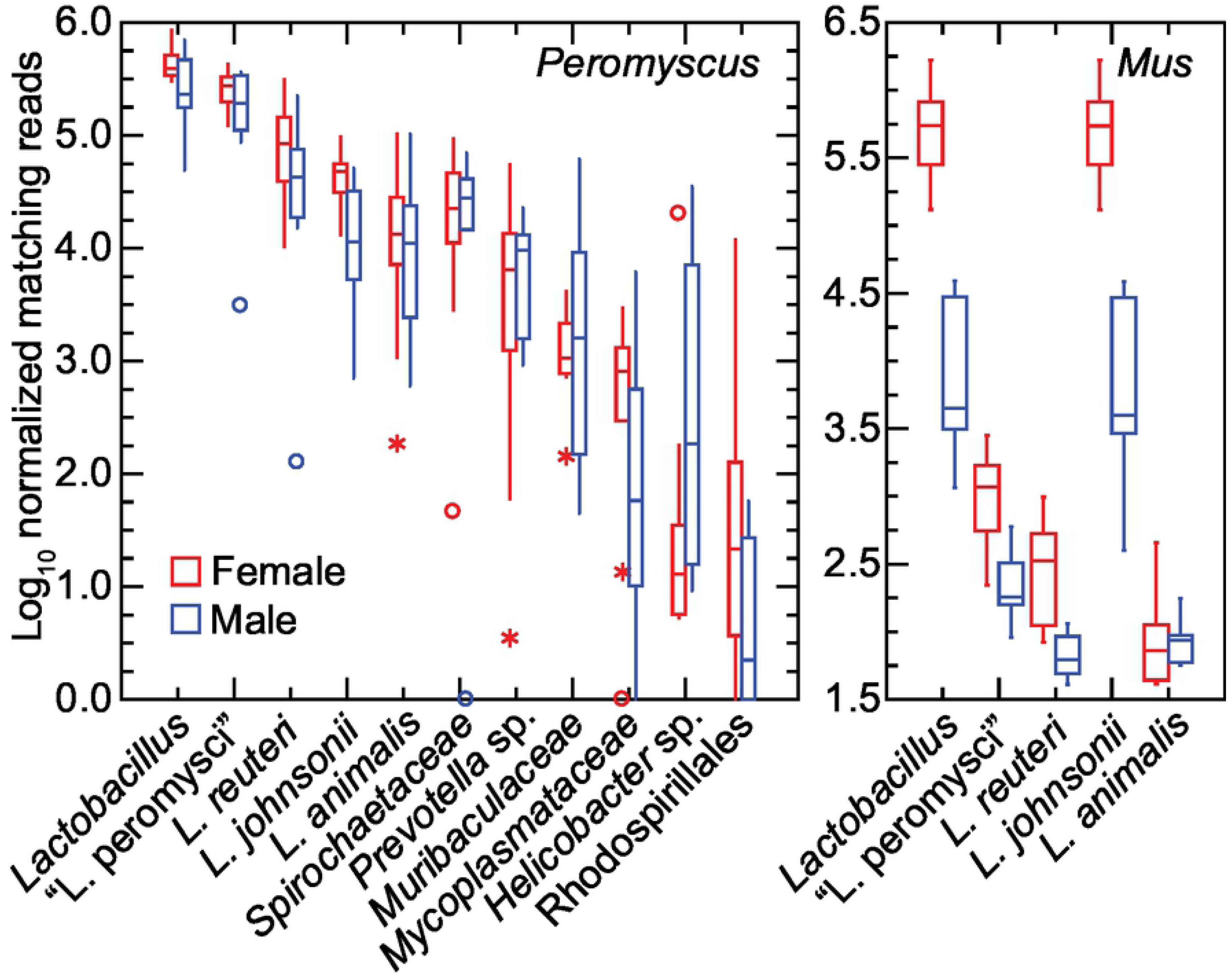
Box-whisker plots of log-transformed normalized reads of gut metagenomes of *P. leucopus* (left panel) and *M. musculus* (right panel) that mapped to chromosomes of *Lactobacillus* spp. or other bacteria by host species and grouped by sex. The references to which reads were mapped were complete chromosomes or partial chromosomes of organisms listed in Table 1. “*Lactobacillus*” in the first position of each panel were the cumulative reads for the four individual *Lactobacillus* species in this analysis.

All four species of the lactobacilli were represented in each of the 20 *P. leucopus* metagenomes. “L. peromysci” and *L. reuteri* tended to be the most common and consistently represented, while *L. johnsonii* and *L. animalis* varied more in prevalences between animals. Other bacteria were also identified in the samples of all or most of the individual animals. The *Spirochaetaceae* bacterium was ∼10-fold less abundant than the cumulative *Lactobacillus* spp. in the *P. leucopus* samples.

The mean number of lactobacilli in aggregate were ∼2-fold more prevalent in *P. leucopus* females than males of the species (*t*-test *p* = 0.03). In *M. musculus* this sex difference for *Lactobacillus* was more pronounced; on average ∼100-fold more reads from female mice mapped to *Lactobacillus* genomes than was found for male mice (*t*-test *p* < 0.001). The differences in amounts of fecal lactobacilli in the sample plausibly account for cluster 3 of the heatmap of Fig 8. *L. johnsonii* largely accounted for these differences between sexes in *M. musculus*; nearly all of the reads mapping to the *Lactobacillus* genus as a whole were mapping to the *L. johnsonii* genome. The three other species identified in *P. leucopus* were either not present or in much lower numbers in this sampling of *M. musculus*. Strains of *L. johnsonii* have been commonly detected in feces of laboratory mice (68).

A limitation to the study was that the LL stock animals were outbred, and the BALB/c mice were inbred. An inbred lineage derived from the LL stock population was not available. On the other hand, this distinction provided a comparison of microbiome diversities between an outbred and inbred population. As expected, there was greater diversity among the outbred samples than the inbred (Fig 10). Another limitation was the dependence on fecal pellets collected at one time point. The samples were from similar age *P. leucopus* and *M. musculus* and were obtained from the animals and then processed on the same day, but for this study we did not assess variation within individuals over time.

**Fig 10.**
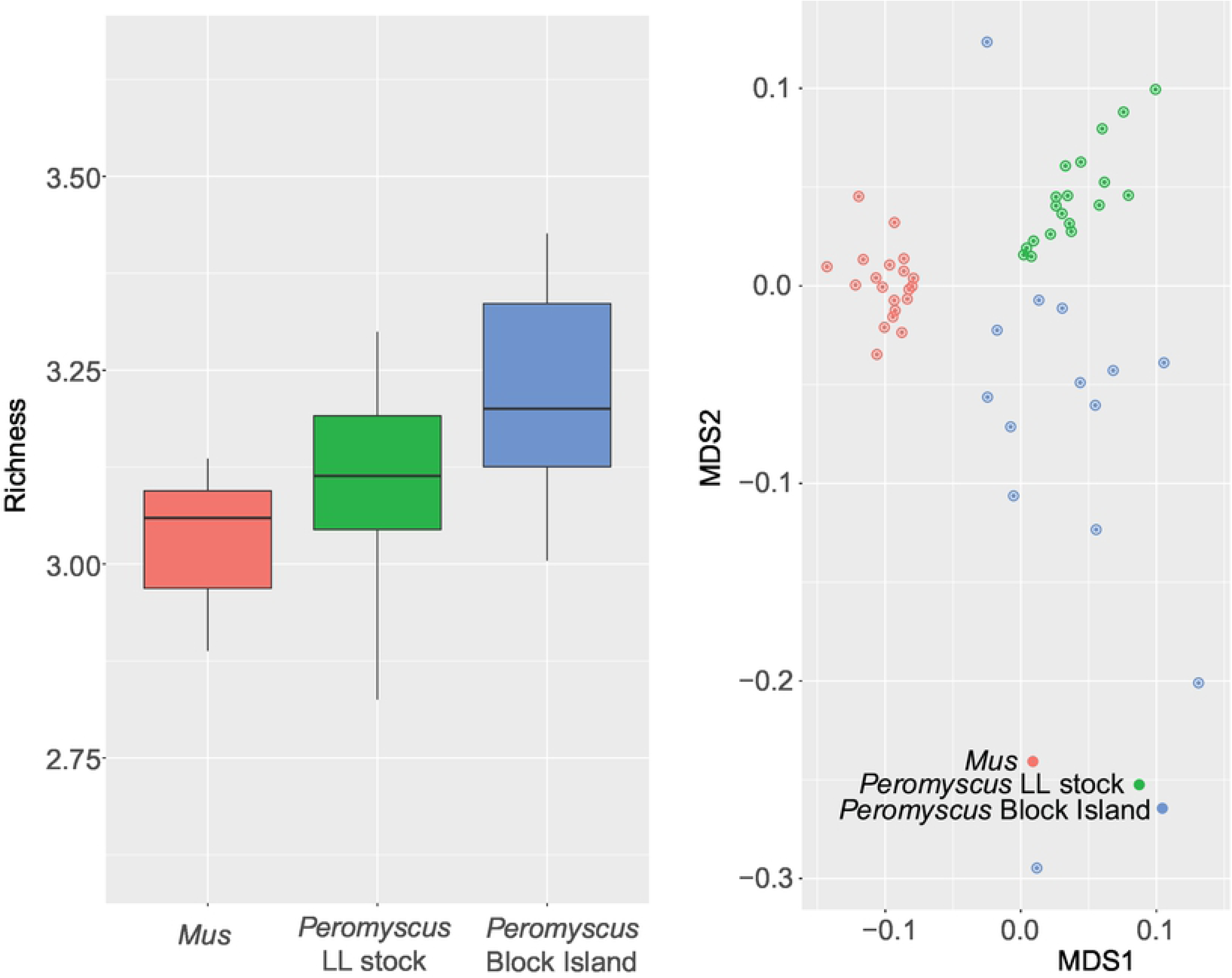
Alpha diversity (left) and beta diversity (right) of gut metagenomes of outbred *Peromyscus leucopus* (green), a natural population of *P. leucopus* (blue), and inbred *Mus musculus* (red). Left panel, box-whisker plots of Shannon’s Index for 20 BALB/c *M. musculus*, 20 LL stock colony *P. leucopus*, and 18 *P. leucopus* trapped on Block Island, RI. The 3 pairwise, 2-tailed *t*-test *p* values between the groups were ≤ 0.02. Right panel, beta diversity by Bray-Curtis measure visualized by multi-dimensional scaling. The greater scattering of the samples from Block Island animals corresponded to the alpha diversity of this group.

### Lactobacilli of the stomach of *P. leucopus*

The differences between *P. leucopus* and *M. musculus* in the amount and species richness of the lactobacilli in their GI microbiota prompted further investigation of *P. leucopus* using histologic, microbiologic and genomic approaches. Fig 11 shows the gross morphology and histology of the stomach of representative LL stock *P. leucopus* animals (69). The difference between forestomach with its stratified squamous epithelium and the discrete region lined with glandular mucosa are indicated in the dissecting scope and higher magnification light microscope views.

**Fig 11.**
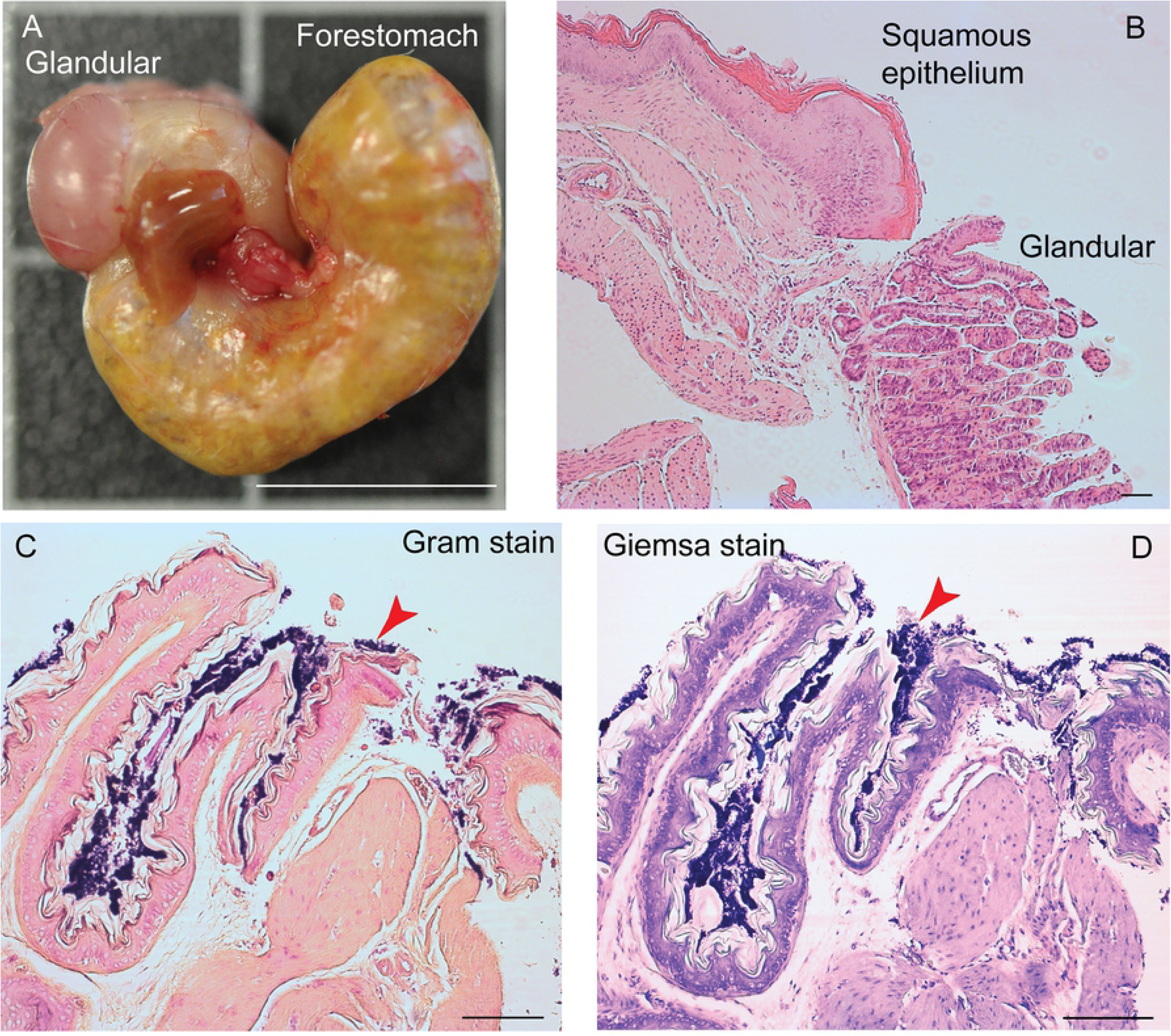
Gross morphology and histology of the stomach of *Peromyscus leucopus* LL stock. The glandular mucosa portions of the stomach and the forestomach with stratified squamous epithelium are indicated. Panel A, whole stomach after dissection. Portions of the esophagus and small intestine are juxtaposed in the center in this view. Bar, 1 cm. Panel B, histology of hematoxylin and eosin-stained section of junction of glandular and squamous epithelium parts. Bar, 100 µm. Panels C and D, Gram stain (C) and Wright-Giemsa stain (D) of sections of squamous epithelium. Bar, 100 µm. Red arrowheads indicate gram-positive bacteria in a biofilm.

Staining of the sections of the fixed gastric tissue with Gram stain or Giemsa stain show a thick layer of gram-positive bacteria on the non-secretory epithelium portion of the stomach. This is similar to Savage et al. noted in the forestomachs of *M. musculus* (70). The appearance is also consistent with the *Lactobacillus* biofilm that was described by Wesney et al. (71).

Two of the species, “L. peromysci” and *L. animalis*, could reliably be distinguished by their distinctive colony morphologies from the isolated strains of *L. reuteri* and *L. johnsonii*, which had colonies of similar appearance (Fig 12). The rough-surfaced, ropy colonies of “L. peromysci” and the compact smooth colonies of *L. animalis* were similar to what Dubos and colleagues described in their study of lactobacilli of the mouse stomach and gut (72).

**Fig 12.**
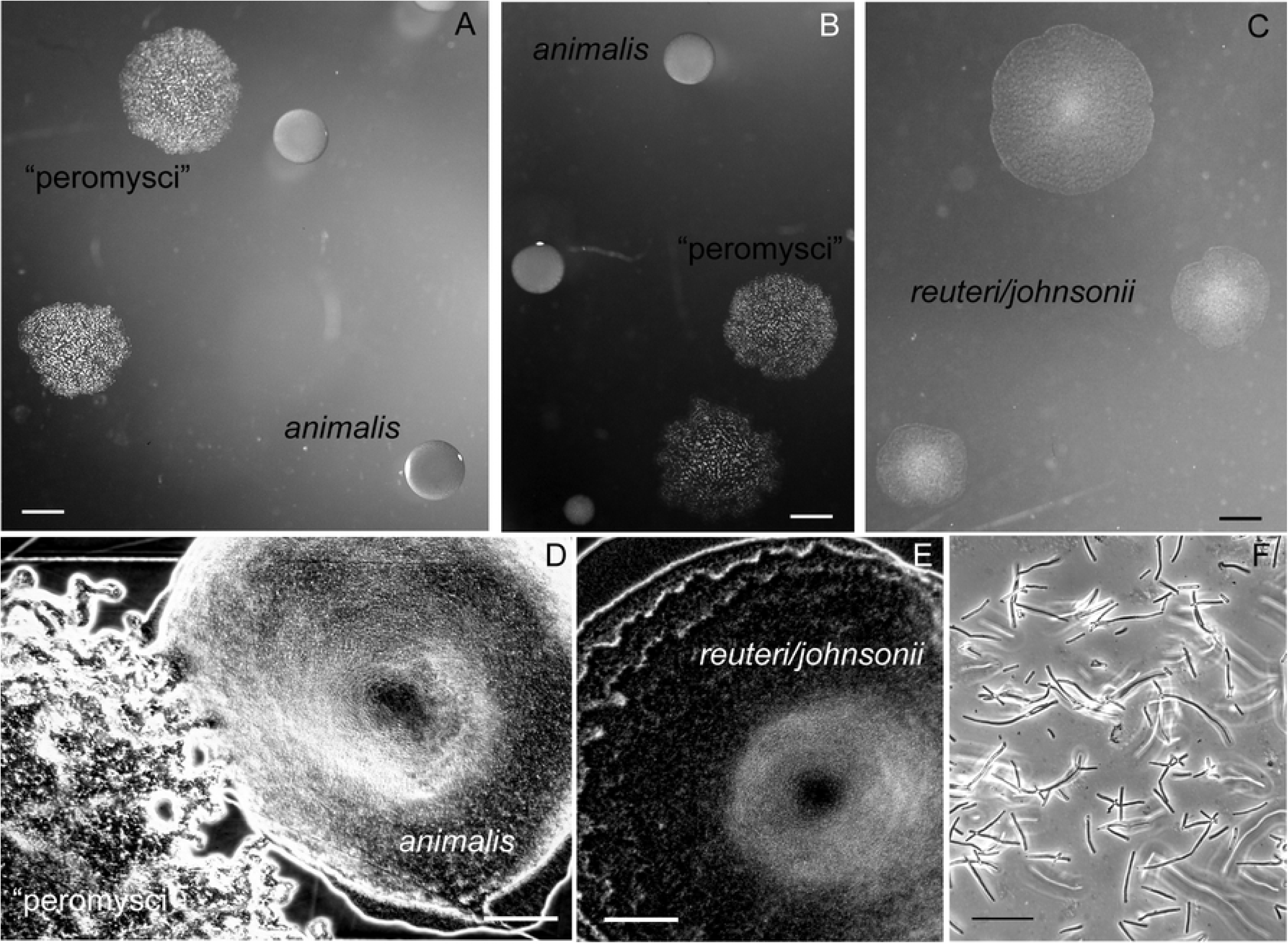
Colonies and cells of lactobacilli of the *P. leucopus* stomach and gut. Panels A-C show representative sizes and morphologies of colonies of “L. peromysci”, *L. animalis*, and the less distinguishable *L. reuteri* and *L. johnsonii*. Bars, 1 mm. Panels D and E show magnified view of colonies of “L. peromysci” and *L. animalis* (D) and that of *L. reuteri* and *L. johnsonii* (E). Bar, 100 µm. Panel F, phase microscopy of wet mount of unconcentrated broth culture of “L. peromysci”. Bar, 10 µm.

We next used a different set of 20 animals of the LL stock, 6 (2 females and 4 males) of which were born at the PGSC facility and 14 (7 females and 7 males) of which were born at U.C. Irvine. All animals were housed at U.C. Irvine for at least 26 weeks before euthanasia, dissection, and cultivation of the stomach tissue and contents.

Mean (95% CI) colony forming units of lactobacilli per gram of stomach tissue on selective medium plates were ten-fold higher in females at 7.4 (1.1-47.4) x 10^9^ than in males at 0.76 (0.40-1.4) x 10^9^ (*t*-test *p* = 0.02 for log-transformed values) (Table 3). There was no discernible association with place of birht, and there was no difference between females and males in the proportions of the lactobacilli were identified as “L. peromysci”, *L. animalis*, and *L. reuteri*/*L. johnsonii*. For five animals, whose lactobacilli were subjected to 16S ribosomal RNA gene PCR and sequencing for confirmation, the *L. reuteri/L. johnsonii*-type colonies are predominantly *L. reuteri*. But *L. johnsonii* was confirmed to be present as well and outnumbered *L. reuteri* in one animal. The results for 3 animals that had been on a 9% fat content diet, which was part of the breeding program, instead of 6% fat content were not distinguishable from those for the other 17.

**Table 3.** Colony forming units (cfu) of *Lactobacillus* spp. in *Peromyscus leucopus* stomach

|  |  |  |  |  |  |
| --- | --- | --- | --- | --- | --- |
| Animal ID | Sex | Log<sub>10</sub> total cfu/gm | % colony type |  |  |
|  |  |  | *animalis* | "peromysci" | *reuteri/johnsonii* |
| 22403 | F | 10.9 | 3 | 32 | 65 |
| 22404 | F | 11.3 | 61 | 4 | 35 |
| 25053 | F | 10.7 | 37 | 60 | 3 |
| 25054 | F | 11.0 | 96 | 4 | <1<sup>a</sup> |
| 25055 | F | 10.4 | 94 | 6 | <1 |
| 25065 | F | 8.9 | 50 | 19 | 31 |
| 25062 | F | 8.5 | 60 | 14 | 26 (26/0)<sup>b</sup> |
| 25063 | F | 8.8 | 52 | 12 | 36 (33/3) |
| 25058 | F | 8.3 | 82 | 14 | 5 (5/0) |
| 22401 | M | 8.6 | 33 | 19 | 48 |
| 22375 | M | 8.8 | 58 | 3 | 40 |
| 22420 | M | 9.7 | 48 | 19 | 33 |
| 22377 | M | 9.0 | 81 | 10 | 9 |
| 26050 | M | 8.4 | 87 | 6 | 7 |
| 25056 | M | 9.6 | 60 | 15 | 26 |
| 25010 | M | 9.0 | 84 | 16 | <1 |
| 25060 | M | 8.8 | 55 | 23 | 23 |
| 25061 | M | 8.3 | 63 | 38 | <1 |
| 25059 | M | 8.5 | 50 | 9 | 41 (36/5) |
| 25011 | M | 9.0 | 61 | 24 | 14 (4/10) |
| Mean (95% CI) | n.a. | 9.3 (8.9-9.8) | 61 (51-70) | 17 (11-23) | ≤ 22 |

<sup>a</sup> <1, below limit of detection by serial dilution on plates
<sup>b</sup> ( / ), % *reuteri* / % *johnsonii* by PCR and 16S ribosomal rDNA sequencing

A separate group of 9 adult LL stock *P. leucopus* (5 females and 4 males) were euthanized after withholding food overnight, and the freshly-excised stomachs were subjected to DNA extraction without prior washing of the stomach. A mean (95% CI) of 477,688 (408,988-546,388) PE250 Illumina reads were obtained for the 9 samples (Table S7 of Supplementary information). These were mapped to the four *Lactobacillus* genomes as references, as described above, as well as to partial chromosomes for *Prevotella* sp. LL70 and *Helicobacter* sp. LL4 (Table 1). For an estimate of the number of mammalian nuclei represented in the stomach extract the *P. leucopus* genome (accession NMRJ00000000.1) served as the reference. Fig 13 shows the distributions of normalized reads mapping to the references as well as to the *P. leucopus* genome and cumulatively to all *Lactobacillus* spp. Females and males were similar by these measures for all these groups. For this group of animals and this analysis, we confirmed the high prevalence of “L. peromysci” in the stomach as well as the comparatively greater representation of *L. reuteri* over *L. johnsonii*. In this sample *L. animalis* was more variable in numbers between animals. As further evidence that the *Helicobacter* sp. was of the enterohepatic type, it was near undetectable in the stomach extract, while a typically abundnant genus in the intestine, *Prevotella*, was present in small numbers in some samples. The lactobacilli in the stomach were about as numerous as the stomach tissue cells constituting the sample.

**Fig 13.**
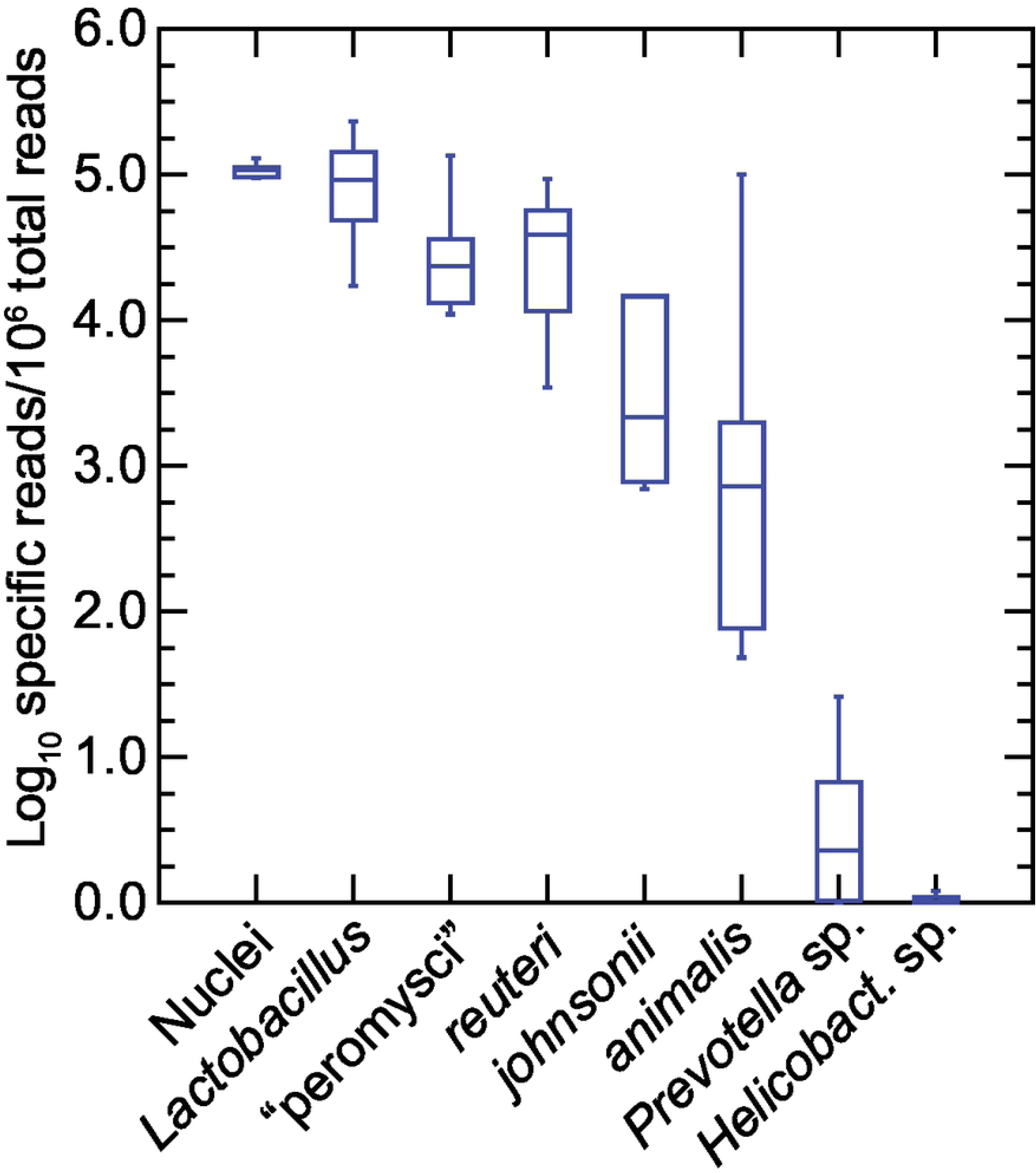
Box-whisker plots of log-transformed normalized reads of total metagenomes of the stomachs of 9 *P. leucopus* (left panel) and *M. musculus* (right panel) that mapped to chromosomes of *Lactobacillus* spp. or other bacteria by host species and by sex. The references to which reads were mapped were complete chromosomes or partial chromosomes of organisms listed in Table 1. “*Lactobacillus*” in the first position of each panel were the cumulative reads for the four individual *Lactobacillus* species in this analysis.

A strain of *L. reuteri* was shown to be the source of biofilm in the GI tract of mice in one study (73), and *L. murinus*, the sister taxa of *L. animalis* (Fig 3), accounted for the biofilm in another study of the upper GI tract of *M. musculus* (74). *L. johnsonii* has also been demonstrated to produce an exopolysaccharide biofilm (75). In a study of germ-free mice in which bacteria were experimentally introduced, *L. taiwanensis*, which is in the same cluster as *L. johnsonii* and *L. gasseri* (67). formed a mixed-species biofilm with *L. reuteri* (76).

One limitation of this experiment is that we may have overlooked species that were not identified because they were not cultivable by our method and conditions, which were microaerophilic, not strictly anaerobic. That said, if cells of such non-cultivable lactobacilli had been present in the feces or stomach, their numbers did not reach a threshold for assembly into contigs of the de novo assembly of the high coverage sequencing and then detection.

### Gut metagenomes of a natural population

The foregoing studies were of animals born and reared under controlled conditions, including the same diet and environmental parameters for all individuals in the group. Infectious diseases and predators were not a variable. The LL stock *P. leucopus* were outbred but the effective population size was small compared to a wild population (18). Which of our findings would hold for animals sampled in their native habitats?

This particular study of a natural population had two specific purposes. The first was to assess the species richness or alpha diversity of microbiota within a given animal and differences in species composition or beta diversity between animals. The second objective’s question of whether any of the *Lactobacillus* species we identified in the stock colony were present in natural populations. For this survey we used fecal pellets from *P. leucopus* that were individually captured and then released on Block Island, several miles off-shore from the North American mainland. At time of capture the animals were identified as to species, sex, and stage of maturity.

We analyzed the data from fecal pellets of 18 different animals (10 females and 8 males), the majority of which were adults, collected from *P. leucopus* captured at different locations on Block Island (Table S14 of Supplementary information). As expected, there was greater variation between individual animals than was observed with the stock colony animals maintained under same conditions. Fig 10 compares the alpha diversity by Shannon index and beta diversity by Bray-Curtis dissimilarity of the inbred BALB/c *M. musculus*, outbred LL stock *P. leucopus*, and the natural population of *P. leucopus* of Block Island.

By algorithmic assignment of reads to taxonomic family, *Lactobacillaceae* was one of the most prevalent bacteria with a mean of ∼5% of reads, but this was over a range of 0.3% to 20%. As was the case for the stock colony *P. leucopus*, the frequency of *Helicobacteraceae* varied more widely between sampled animals than for comparably-prevalent taxa: a mean of ∼1% but ranging from 0.03% to 12%. The frequency of a parabasalid protist, by the criterion of *Trichomonadidae* reads, in the metagenomes was similar to what we observed in the metagenome of the stock colony *P. leucopus*: the mean was 0.11% with a range of 0.02 to 0.62%. This was an indication that the *T. muris* was autochthonous in *P. leucopus*, but we did not have direct observation of the protozoa to confirm that.

Using the chromosome sequences of the four *Lactobacillus* species and partial chromosome sequences of *Spirochaetaceae* bacterium LL50, *Prevotella* sp. LL70, and *Helicobacter* sp. LL4 as references, we mapped and counted reads, as described for the LL stock and *M. musculus* study above. Fig 14 summarizes results for the 18 animals grouped by sex. Lactobacilli were common but, as seen with family level matching, there was greater variation between samples of the different animals than was observed for colony animals. There was also substantial variation in prevalences of the *Spirochaetaceae* bacterium and the *Prevotella* species. In most of the samples there was scant evidence of the *Helicobacte*r species but in two animals, there were higher numbers of this organism, reaching 7% of the total reads in one fecal sample.

**Fig 14.**
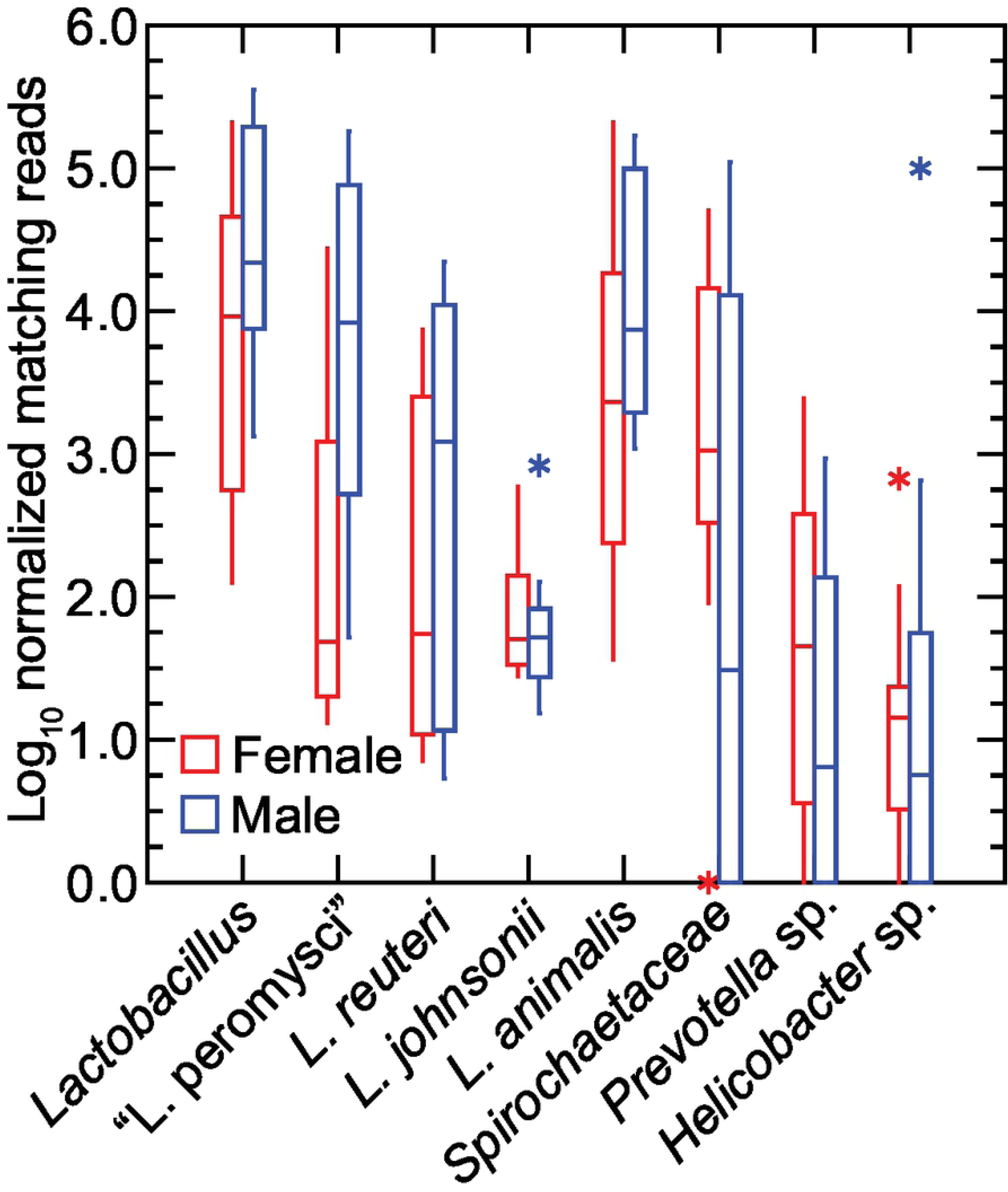
Box-whisker plots of log-transformed normalized reads of fecal metagenomes of 5 female (red) and 4 male (blue) *P. leucopus* of a natural population of Block Island, Rhode Island. The reference genomes and other sequences were those described for Fig 13 and in addition the partial chromosome sequence of *Spirochaetaceae* sp. LL50. As an estimate of the number of mammalian cells in the extract, “nuclei” corresponded with normalized reads mapped to *P. leucopus* genome.

Among the lactobacilli used as references for this analysis, the two most prevalent species were “L. peromysci” and *L. animalis*. *L. reuteri* overall was about 10-fold lower in frequency, and *L. johnsonii* was about a hundred-fold lower in frequency. It is likely that reads called as *L. johnsonii* were the result of complete or partial matching to chromosomal loci that were highly conserved across the genus. Unlike the stock colony *P. leucopus* and the *M. musculus*, in this sampling of the Block Island population the samples from female animals had marginally lower representation of lactobacilli in the fecal samples than males.

To better characterize the two predominant *Lactobacillus* species in this set, we assembled contigs of reads mapping to “L. peromysci” or *L. animalis* from a higher coverage sequencing of the DNA of one of the Block Island samples. This yielded 51 of the 53 genes for ribosomal protein genes for a strain of *L animalis*, which was designated 7442BI, and several core or housekeeping genes for the “L. peromysci”-like organism, which was designated BI7442 (Table 1). Fig 15 shows phylograms of DNA sequences for these and related *Lactobacillus* species or strains. The concatenated sequence of the BI7442 isolate was 99.2% identical over the 11,252 nt aligned with the corresponding sequences of the LL6 isolate of “L. peromysci”. Isolate 7442BI was comparatively more distant from the LL1 isolate of *L. animalis* in the stock colony but still clustered with it rather than with other examples of *L. animalis*.

**Fig 15.**
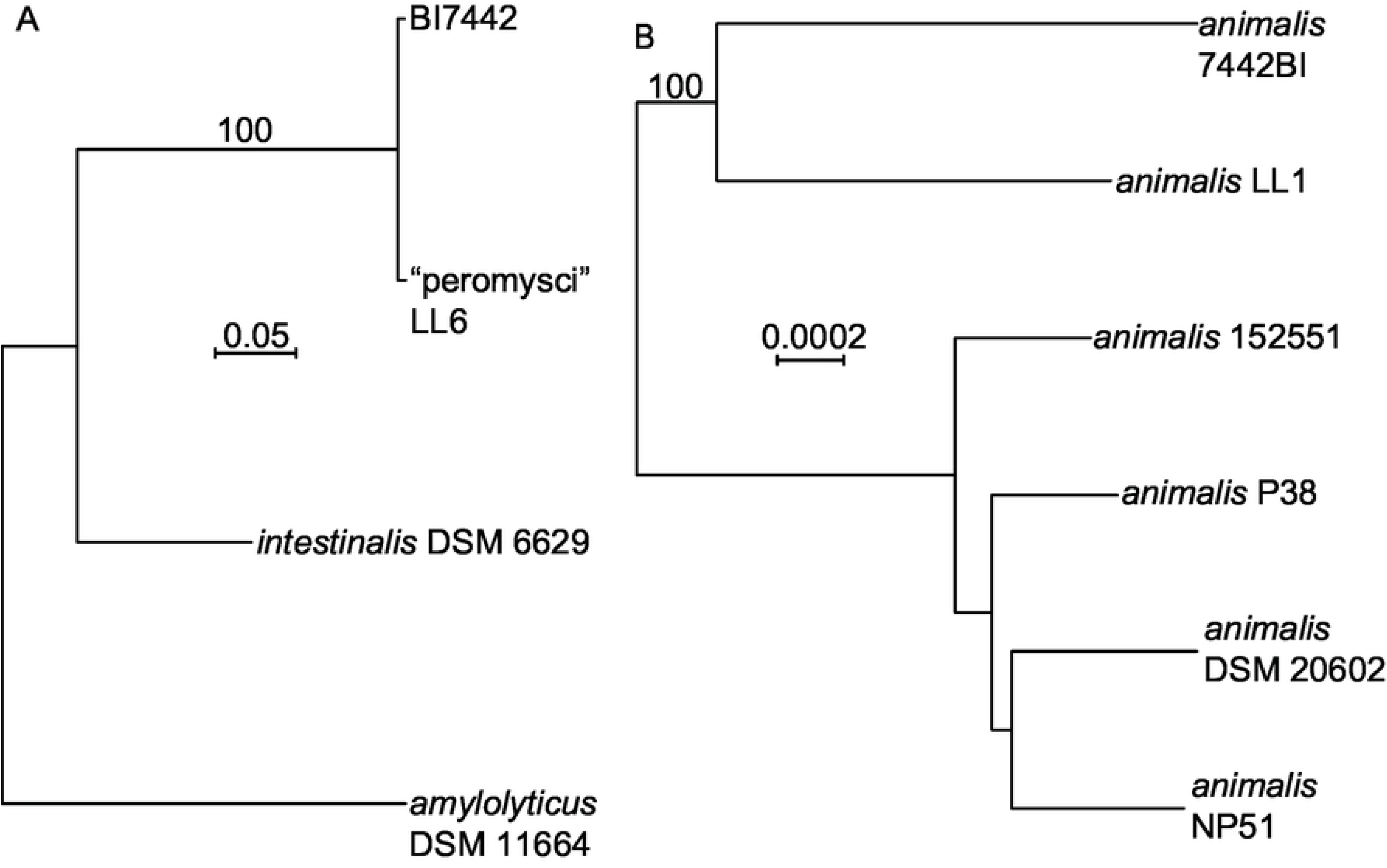
Distance phylograms of concatenated codon-aligned nucleotide sequences of two *Lactobacillus* spp. of *P. leucopus* of Block Island, Rhode Island. Panel A, 10,152 positions of *ftsK, ftsZ, dnaA, dnaN, ileS*, and *topA* of “L. peromysci” LL6 and BI7442 and two other *Lactobacillus* species. Panel B, 18,552 positions of 51 of 53 ribosomal protein genes of six *L. animalis* strains, including LL1 and 7442BI. The distance criteria were Jukes-Cantor. The scales for distance are shown in each panel. Percent bootstrap (100 iterations) support values of ≥ 80% at a node are shown.

The sample size was limited, and we did not attempt culture isolations from the pellets. But the source of samples from *P. leucopus* was notable for its location on an island where *B. burgdorferi* is enzootic (77) and the risk of infection for residents and visitors is high (78). If there were to be future interventions targeting *P. leucopus* to interrupt disease transmission, Block Island would likely be a candidate site for this application.

This survey also documented that a strain or strains of “L. peromysci” and *L. animalis* are present in the native deermice. The high degree of sequence identity between two “L. peromysci” examples, whose origins were North Carolina and a New England island, long separated from the mainland, suggest that this newly-discovered species is autochthonous and plausibly a narrowly host-restricted symbiont of *P. leucopus*. Host-range restrictions of lactobacilli for the stomachs of mice were demonstrated by Wesney and Tannock (71). Supporting an assignment of a symbiont lifestyle was its smaller genome size and lower % GC content of this species in comparison with *L. reuteri* and *L. johnsonii* with their broader host ranges (79).

## Conclusions

Six decades ago René Dubos (of the epigraph), Russell Schaedler, and their colleagues at what is now Rockefeller University reported in a series of ground-breaking papers on the “fecal flora” of mice and variations in that microflora between mouse strains (80–82). They associated differences in gastrointestinal flora with growth rates of the mice and the mouse’s susceptibility to infection and endotoxin. A featured group of bacteria in their studies were lactobacilli. They showed that this group of bacteria were present in large numbers in the feces and that they predominated (up to 10^9^ cfu per g of homogenate) in the stomachs of the mice (70), similarly to what we observed in *P. leucopus*. As their studies first intimated, the rodent may plausibly owe as much to the genomes of their microbiota as to the nuclei and mitochondria of their somatic cells for either ameliorating or exasperating disease (83).

There are also implications of our findings for development of oral vaccines that target *P. leucopus* to block transmission of pathogens either from tick to the reservoir or from the reservoir to the tick. Two of the candidate vehicles for the bait delivery of recombinant vaccine antigens to rodents have been *E. coli* and a *Lactobacillus* species (30, 84). In neither case were the strains known to be adapted for life in *P. leucopus*. Success rate for achieving a protective response may be enhanced by use as the bacterial vehicle microbes that are adapted to *P. leucopus*. Such organisms presumably would more likely than an allochthonous bacterium to stably colonize and then proliferate to numbers large enough for the recombinant protein to elicit the sought-after immune responses.

Finally, this exploration and curation of microbes in the gut of the white-footed deermouse concludes the third leg of our project on the total genome of representative animals of the species: the nuclear genome (18), the mitochondrial genome (17), and now the GI microbiome (85). This provides a foundation for testing of hypotheses by selective manipulation of the microbiota, for instance, by specifically targeting a certain species with a lytic phage or bacteriocin, to which it is not immune, and then evaluating the phenotype of the animal after this “knock-out”. Now that there is an annotated *P. leucopus* genome with millions of SNPs identified (UC Santa Cruz genome browser; http://googl/LwHDr5) it also feasible to investigate through forward genetics the host determinants of particular bacterial associations and for which there is evidence of variation within a population. An example would be the *Helicobacter* species that was highly variable in prevalence in both the wild animals and the stock colony animals.

## Materials and methods

### Colony animals

This study was carried out in strict accordance with the recommendations in the Guide for the Care and Use of Laboratory Animals of the National Institutes of Health. At the University of California Irvine protocol AUP-18-020 was approved by the Institutional Animal Care and Use Committee (IACUC)-approved protocol. Adult outbred *P. leucopus* of the LL stock were purchased from the Peromyscus Genetic Stock Center (PGSC) of the University of South Carolina (86). The closed colony of the LL stock was founded with 38 animals captured near Linville, NC in the mid-1980’s. Some of the LL stock animals in the study were bred at the University of California Irvine’s animal care facility from pairs originating at the PGSC. Adult BALB/cAnNCrl (BALB/c) *M. musculus* were purchased from Charles River. For the species comparison experiment both the PGSC-bred *P. leucopus* and *M. musculus* animals were housed in Techniplast individual ventilated cages in vivarium rooms with a 12 hour-12 hours light-dark cycle, an ambient temperature of 22 ± 1 °C, stable humidity, and on an ad libitum diet of 2020X Teklad global soy protein-free extruded rodent chow with 6% fat content (Envigo, Placentia, CA). Other animals of PGSC origin, including for the high-coverage gut metagenome study, were also housed under the same conditions and on the same diet. Twenty U.C. Irvine-bred animals were under the housing conditions and on the diet except for three (1 female and 2 males) that were on 2019 Teklad global protein extruded rodent chow with 9% fat content. Before euthanasia with carbon dioxide asphyxiation and cervical dislocation and then dissection of the stomach, food but not water was withheld for 12 h for selected animals. *P. leucopus* studied at the PGSC were under IACUC-approved protocol 2349-101211-041917 of the University of South Carolina and were euthanized by isoflurane inhalation.

### Field site and animal trapping

The study was performed under IACUC-approved protocol AC-AAAS6470 of Columbia University (77). Block Island, located 23 km from mainland Rhode Island, is part of the Outer Lands archipelagic region, which extends from Cape Cod, MA through to Staten Island, NY. Block Island is 25.2 sq. km, about 40% of which is maintained under natural conditions. The agent of Lyme disease *Borreliella burgdorferi* is enzootic on the island (87) Animals were trapped at three locations: 1, a nature conservation area (41.15694, −71.58972); 2, private land with woodlots and fields (41.16333, −71.56611); and 3, Block Island National Wildlife Refuge (41.21055, −71.57222). Trapping was carried out during the May-August period with Sherman live traps (H.B. Sherman Traps, Inc. Tallahassee, FL) that were baited with peanut butter, oats, and sunflower seeds. Traps were arranged in nine 200 m transects with one trap placed every 10 m for a total of 180 traps at each location. Animals were removed from traps, weighed, sexed, and assessed as to age (adult, subadult, or juvenile) by pelage. Fecal pellets were collected and kept at −20 °C on site, during shipment and until DNA extraction. The species identification of the source of the fecal pellets as *P. leucopus* was confirmed by sequencing of the D-loop of the mitochondrion as described (17).

### Cultivation and enumeration of bacteria

Lactobacilli were initially isolated and then propagated on Rogosa SL agar plates (Sigma-Aldrich) in candle jars at 37° C. Gram-negative bacteria and specifically *Escherichia coli* were isolated and propagated on MacConkey Agar plates (Remel) incubated in ambient air at 37° C. For determinations of colony forming units (cfu) homogenates of stomach, cecum, or fecal pellets were suspended and the serially diluted in phosphate-buffer saline, pH 7.4, before plating in 100 µl volumes on solid media in 150 mm x 15 mm polystyrene Petri dishes. Colonies were counted manually. Liquid cultures of *Lactobacillus* spp. isolates or *E. coli* were in Difco Lactobacilli MRS Broth (Becton-Dickinson) or LB broth, respectively, and incubated at 37° C on a shaker. Bacteria were harvested by centrifugation at 8000 x g for 10 min. Antibiotic susceptibilities were determined by standard disk testing on Mueller-Hinton Agar (Sigma-Aldrich) plates and ciprofloxacin 5 µg, gentamicin 10 µg, ampicillin 10 µg, and sulfamethoxazole 23.75 – trimethoprim 1.25 µg BBL Sensi-Disc antibiotic disks (Becton-Dickinson) according the manufacturer instructions.

### Histology

After the stomachs were removed from two euthanized *P. leucopus* LL stock adult females, they were opened longitudinally, gently flushed with PBS, and fixed in 10% buffered formalin (Thermo Fisher Scientific). Histological and histochemical analysis was performed on paraffin block sections of the stomach with Hematoxylin and Eosin, Wright-Giemsa and Gram stains (Abcam, Cambridge, UK).

### Microscopy, photography, and video

Photographs of colonies on plates were taken with a Nikon Df DSLR camera and 60 mm Nikkor AF-S Micro lens with illumination by incident light above and reflected light below the plates on an Olympus SZ40 dissecting scope. An Olympus BX60 microscope with attached QIClick CCD camera and Q-Capture Pro7 software (Teledyne Photometrics, Tucson, AZ) was used for low-magnification images of colonies under bright light microscopy and 400X images under phase and differential interference microscopy. Histology slides were examined on a Leica DM 2500 microscope equipped with a MC120 HD digital camera (Leica Microsystems, Buffalo Grove, IL).

### DNA extractions

DNA from fresh and frozen fecal pellets, from tissue of stomach and cecum, and from bacteria harvested from broth cultures were extracted with ZymoBIOMICS™ DNA Miniprep or Microprep kits (Zymo Research, Irvine, CA). Freshly-dissected, unwashed tissues were cut into small pieces before trituration and then homogenization in the lysis buffer. DNA concentration was determined by NanoDrop spectrophotometer and Qubit fluorometer (Thermo Fisher Scientific).

### PCR

The near-complete 16S ribosomal RNA gene for Lactobacillus spp. was amplified using PCR using custom primers for the genus *Lactobacillus*: forward 5’-CCTAATACATGCAAGTCG and reverse 5’-GGTTCTCCTACGGCTA. The Platinum Taq polymerase and master mix (ThermoFisher Scientific) contained uracil-DNA glycosylase. On a T100 thermal cycler (BioRad) PCR conditions (°C for temperature) The conditions were 37° for 10 min, 94° for min, 40 cycles of 94° for 10 s, 55° for 30 s, and 72° for 45 s. The 1.5 kb PCR product was isolated and purified from agarose gel using the NucleoSpin Gel and PCR Clean-up kit (Takara). The product was subjected to Sanger dideoxy sequencing at GENEWIZ (San Diego, CA).

### Whole genome sequencing, assembly, and annotation

Long reads were obtained using an Oxford Nanopore Technology MinION Mk1B instrument with Ligation Sequencing Kit, R9.4.1 flow cell, MinKnow v. 19.6.8 for primary data acquisition, and Guppy v. 3.2.4 for base calling with default settings. Paired-end short reads were obtained on a MiSeq sequencer with paired-end v2 Micro chemistry and 150 cycles (Illumina, San Diego, CA). The library was constructed using the NEXTflex Rapid DNA-Seq kit (Bioo Scientific, Austin, TX), the quality of sequencing reads was analyzed using FastQC (Babraham Bioinformatics), and reads were trimmed of Phred scores <15 and corrected for poor-quality bases using Trimmomatic (88). A hybrid assembly was carried out with Unicycler v.0.4.7 (89) with default settings and 16 threads on the High Performance Computing cluster of the University of California Irvine. Assembly of short reads alone were performed with the Assembly Cell program of CLC Genomics Workbench v. 11 (Qiagen). Annotation was provided by the NCBI Prokaryotic Genome Annotation Pipeline (90). Putative bacteriocins and their associated transport and immunity functions were identified by BAGEL4 (91, 92). For other analyses paired-end reads were mapped with a length fraction of 0.7 and similarity fraction of 0.9 to whole genomes sequences or concatenated large contigs representing partial genomes (Table 1). Mapped reads were normalized for length of reference sequence and for total reads after quality control and removal of adapters.

### Metagenome sequencing

The library was constructed using NEXTflex Rapid DNA-Seq kit v2 (Bioo Scientific) and the NEXTflex Illumina DNA barcodes after shearing the DNA with a Covaris S220 instrument, end repair and adenylation, and clean-up of the reaction mixture with NEXTFLEX Clean Up magnetic beads (Beckman Coulter, Brea, CA). The library was quantified by qPCR with the Kapa Sybr Fast Universal kit (Kapa Biosystems, Woburn, MA), and the library size was determined by analysis using the Bioanalyzer 2100 DNA High Sensitivity Chip (Agilent Technologies). Multiplexed libraries were loaded on either an Illumina HiSeq 2500 sequencer (Illumina, San Diego, CA) with paired-end chemistry for 250 cycles or a MiSeq Sequencer (Illumina, San Diego, CA) with paired-end v2 Micro chemistry and 150 cycles. The Illumina real time analysis software RTA 1.18.54 converted the images into intensities and base calls. De novo assemblies were performed with De Novo Assembly v. 1.4 of CLC Genomics Workbench v. 11 with the following settings: mismatch, insertion, and deletion costs of 3 each; length fraction of 0.3, and similarity fraction of 0.93.

### 16S ribosomal RNA analysis

The same DNA extract used for the metagenome sequencing at University of California Irvine was submitted to ID Genomics, Inc. (Seattle, WA) and subjected the company’s 16S rRNA Metagenomics service (http://idgenomics.com/our-services), which used the 16S Metagenomics v. 1.01 program (Illumina). Of the 333,358 reads 82%, were classified as to taxonomic family.

### Metagenome analysis

Fastq files were uploaded to the metagenomic analysis server MG-RAST (https://www.mg-rast.org) (93). Reads were joined using join paired reads function on the browser and filtered for *Mus musculus* v37 genome. Artificial replicate sequences produced by sequencing artifacts were removed by the method of Gomez-Alvarez et al. (94). Low quality reads (Phred score <15 for no more than 5 bases) were removed using SolexQA, a modified DynamicTrim protocol (95). The output of the MG-RAST protocol was analyzed in *R* using the *vegan* package (https://cran.r-project.org, https://github.com/vegandevs/vegan). Alpha diversity was expressed by the Shannon’s Diversity Index, which accounts for evenness as well as richness (96). Beta diversity expressed as the Bray-Curtis Dissimilarity statistic (97) was calculated using the *avgdist* function with 1000 sample depth, the median as the function, and 100 iterations (https://github.com/vegandevs/vegan/blob/master/man/avgdist.Rd). Data was visualized using non-metric multidimensional scaling in two dimensions (98). MicrobiomeAnalyst (https://www.microbiomeanalyst.ca) (99) was used for hierarchical clustering by distance criterion and by means of Pearson correlations. The DFAST prokaryotic genome annotation pipeline (https://dfast.nig.ac.jp) was used for annotation of incomplete chromosomes and large contigs (100). For *Lactobacillus* spp. and the *Helicobacter* sp. the lactic acid bacteria database and *Helicobacter* database, respectively, options were chosen. Alignments and phylogenetic analysis were carried out with the SeaView v. 4 suite (101).

### Data availability

The resources for the several new sequences for genomes, large contigs, and individual genes that described here are listed in Table 1. The accession numbers for the annotated genomes and plasmids of *L. animalis* LL1, *L. reuteri* LL7, and “L. peromysci” LL6 are given in Bassam et al. (24). Fig 2 and its legend provides accession numbers for 16S ribosomal RNA gene sequences of other *Lactobacillus* species and strains. The nucleotide sequences of ribosomal proteins for different species and strains of *Lactobacillus* and *Helicobacter* were obtained from the Ribosomal MLST database of PubMLST (https://pubmlst.org/rmlst/) and given in Table S3 of Supplementary information. Additional accession numbers for individual genes of other organisms as references are given in Table S5 of Supplementary information. Hyperlinks to the long (Nanopore) and short (Illumina) sequence reads at the Sequence Read Archive or MG-RAST database are provided in Table 1 and Tables S7 and S13 of Supplementary information.

### Statistical analysis

Normalized reads and other values whose distributions spanned more than one order of magnitude were log-transformed before parametric analysis by 2-tailed *t*-test. Inverse transformation was carried out to provide nonparametric means and corresponding asymmetric 95% confidence intervals. The *Z*-score was the number of standard deviations below or above the population mean a give raw value was. The False Discovery Rate (FDR) with corrected *p* value was estimated by the method of Benjamini and Hochberg (102). The box-whisker plot graphs were made with SYSTAT v. 13.1 software (Systat Software, Inc.).

## Acknowledgements

We thank Emma Keay and Vanessa Cook at the University of California Irvine and Asieh Naderi and Vimala Kaza at the University of South Carolina for their contributions to the study. The research was supported by National Institutes of Health grant R21 AI136523 to A.G.B., National Science Foundation grants 1736150 and 1755670 to H. Kiaris (University of South Carolina), and National Science Foundation (Division of Integrative Organismal Systems) grant IOS1755286 to D.M.T. and M.A.D.

## Supporting information

Fig S1. Rarefaction curve for high-coverage sequencing of *Peromyscus leucopus* LL stock gut metagenome.

Fig S2. MG-RAST analysis high-coverage sequencing of *Peromyscus leucopus* LL stock gut metagenome by phylum.

Fig S2. MG-RAST analysis high-coverage sequencing of *Peromyscus leucopus* LL stock gut metagenome by subsystems.

Fig S4. Maximum likelihood phylogram of 470 aligned amino acids of DNA polymerase type B, organellar and viral family protein of *Tritrichomonas* sp. LL5 (Table 1) and homologous proteins of other protozoa (*T. foetus*, *Trichomonas vaginalis*, *Entamoeba invadens*, and *Giardia* sp.), oocytes (*Aphanomyces astaci* and *Thraustothea clavata*), and bacteria (Division WS6 bacterium and *Haliea* sp.). The GenBank accession numbers for the sequences are given next to organism name. The bootstrapped tree was generated with PhyML with the setstings of the LG model, 4 rate classes, and 100 replicates.

Table S1. Comparison of 16S sequence-based and metagenome-based identification of bacterial families in fecal pellets of LL stock *P. leucopus*

Table S2. Metagenome by taxonomic family of fecal pellets of LL stock *P. leucopus*

Table S3. Sources of coding sequences for ribosomal proteins at rMLST database of http://mlst.org

Table S4. Putative bacteriocins and associated transport proteins and immunity proteins of 3 *Lactobacillus* species of *Peromyscus leucopus*

Table S5. Accession numbers of sequences of other species

Table S6. Statistics for gut metagenomes of *Peromyscus leucopus* and *Mus musculus*

Table S7. *Peromyscus leucopus* and *Mus musculus* gut metagenome accession numbers

Table S8. Replicates of libraries of *Peromyscus leucopus* DNA extracts of fecal pellets

Table S9. Log10 of mean number of normalized reads for gut metagenomes of *Peromyscus leucopus* and *Mus musculus* by families with >3000 reads

Table S10. Log10 of reads matched to function for metagenomes of fecal pellets of *Peromyscus leucopus* and *Mus musculus*

Table S11. Map-to-reference normalized PE250 reads (log10) for feces and stomach sample against *Lactobacillus* spp. and selected other bacteria

Table S12. Comparison of gut metagenomes of *Peromyscus leucopus* and *Mus musculus* by KEGG orthology gene

Table S13. Reads of gut metagenomes of *Peromyscus leucopus* and *Mus musculus* by KEGG orthology gene for individual animals and data for analysis of Table S11

Table S14. Block Island *Peromyscus leucopus* fecal metagenome Movie S1. Cecal content with *Tritrichomonas* flagellates

